# Epigenomic profiling of cerebrospinal fluid cells identifies immune regulatory alterations and implicates protocadherins in multiple sclerosis

**DOI:** 10.64898/2026.02.25.708054

**Authors:** Yanan Han, Galina Yurevna Zheleznyakova, Chiara Sorini, Majid Pahlevan Kakhki, Nicolas Ruffin, Heng Liang, Nils Hallen, Chandana Rao Prakash, Viveca Beckers, Elena Ivanova, Mohsen Khademi, Mikael C. I. Karlsson, Fredrik Piehl, Tomas Olsson, Gavin Kelsey, Lara Kular, Maria Needhamsen, Maja Jagodic

## Abstract

Multiple sclerosis (MS) is a chronic inflammatory disease of the central nervous system (CNS), where DNA methylation may play a role by connecting genetic and environmental risk factors. We performed whole-genome DNA methylation profiling of cerebrospinal fluid (CSF) cells from relapsing-remitting MS patients and matched controls, identifying 2,710 differentially methylated positions (DMPs) and 4,330 regions (DMRs). These changes were enriched in immune signaling, adhesion and migration processes, and were accompanied by corresponding RNA expression changes. MS-associated methylation changes enriched in the *cohesin chromatin regulation pathway* mapped to enhancers of T helper 17 (Th17) cells, whereas in other T cell types they were mapping to bivalent enhancers and repressed chromatin. Notably, this pathway comprised multiple Protocadherin (*PCDH*) genes, typically expressed in neuronal cells, that displayed consistent methylation and expression changes in CSF cells. Expression of shared intracellular domain of PCDHγ cluster proteins was confirmed in peripheral blood T cells by flow cytometry as well as expression of *PCDH*γ cluster genes in memory CD4^+^ T cell subsets. Moreover, co-expression analysis suggests a role of *PCDH* genes in aryl hydrocarbon receptor (AHR) signaling. In summary, DNA methylation changes in CSF resident cells reflect dysregulated T cell activation and migration in MS and suggest a novel role of protocadherin molecules in MS pathogenesis.

## Introduction

Multiple sclerosis (MS) is a chronic autoimmune disease of the central nervous system (CNS) affecting approximately 2.8 million individuals worldwide^1, 2^. Initially, disease is typically characterized by recurrent relapses driven by autoimmune-mediated inflammatory attacks on the CNS, followed by periods of remission, referred to as relapsing-remitting MS (RRMS)^3^. Although current immunomodulatory therapies can alleviate symptoms and drastically reduce relapse rates, none provide a cure and their impact on slowing down disease progression remains to be established^4^. Individual’s genetic background contributes to both the susceptibility and severity of MS^5, 6, 7^ while environmental exposures, including Epstein-Barr virus (EBV) infection, vitamin D deficiency, and smoking, further modulate disease risk and progression^8^. Recent compelling evidence suggests that genetic risk variants and EBV infection act synergistically to disrupt immune tolerance, thereby promoting autoreactive T cell responses against CNS antigens^9, 10, 11, 12, 13^. Despite these insights, molecular mechanisms linking the risk factors to pathogenic immune activity remain incompletely understood, limiting our ability to develop more precise therapies.

This has led to increasing interest in epigenetic mechanisms, which may mediate the interaction between genes and environmental exposures and serve as a molecular interface between the risk factors and altered immune cell function in people with MS (pwMS)^14^. DNA methylation, the most extensively studied epigenetic modification, involves the addition of a methyl group to cytosine residues, typically at CpG dinucleotides. When present in promoter or enhancer regions, DNA methylation can inhibit transcription by blocking binding of transcription factors or by recruiting repressive chromatin complexes. DNA methylation has been reported to mediate the effects of genetic and lifestyle factors in MS^14, 15, 16, 17^. Moreover, large-scale and multi-cell-type studies consistently identified the human leukocyte antigen (HLA) region, in particular the strongest MS risk gene, *HLA-DRB1*15:01*, as the most robust region of differential methylation in pwMS^14, 15, 18, 19^. The *HLA-DR* genes encode the major histocompatibility complex (MHC) class II molecules that present antigens to CD4⁺ T cells, strongly implicating the critical role of CD4^+^ T helper (Th) cells in the breakdown of tolerance to self-antigens in pwMS. *HLA-DRB1*15:01* is associated with allele-specific DNA hypomethylation and increased expression in immune cells, linking genotype to altered epigenetic regulation and immune activation^14, 15^. Beyond HLA, DNA methylation also regulates genes encoding lineage-defining transcription factors, such as retinoic acid receptor-related orphan receptor gamma t (*RORC*) and forkhead box P3 (*FOXP3*), shaping the balance between pro-inflammatory Th17 and regulatory (Treg) cells^20, 21, 22, 23^. Disruption of this balance is strongly implicated in MS pathogenesis^24, 25, 26^. However, most methylation studies have focused on peripheral blood, where pathogenic T cell subsets are rare and may not fully reflect compartmentalized CNS responses.

Cells circulating in the cerebrospinal fluid (CSF) provide a unique window into the immune processes occurring within the CNS, as CSF resides in close proximity to demyelinating lesions and reflects active neuroinflammation. CSF cells are predominantly composed of T lymphocytes, with CD4⁺ T cells representing the major population. Comparative profiling of CSF cells and peripheral blood mononuclear cells (PBMC) has shown that CSF cells exhibit stronger pathogenic signatures, with CD4⁺ T cells displaying heightened expression signatures of activation, tissue adaptation, and trafficking in MS CSF^27, 28, 29^. Recent single-cell studies have further highlighted MS-specific immune states within the CSF, revealing broad cellular and transcriptional alterations in neuroinflammation^30^, the presence of CNS-resident immune cells detectable in CSF^31^, T follicular helper (Tfh) cells that can facilitate B cell infiltration into the CNS^32^, and clonally expanded cytotoxic T cell populations in pwMS^33,34^. These cellular states are marked by transcriptional programs characterized by T cell activation, interferon (IFN) and tumor necrosis factor (TNF) signaling and cytotoxicity^33^. Although single-cell transcriptomics has provided important insights, its limited gene coverage restricts the detection of lowly expressed but functionally critical regulators such as transcription factors. Moreover, transcriptomic profiles offer only a cross-sectional snapshot of cellular states at the time of sampling, in contrast to epigenetic modifications that can capture more stable or historical regulatory imprints.

In this study, we sought to elucidate molecular mechanisms underlying MS immunopathology by identifying and functionally characterizing genome-wide DNA methylation changes in CSF cells, which are enriched for pathogenic immune activity. A sequencing approach optimized for low-input samples^35, 36^ enabled us to generate the first comprehensive methylome of CSF infiltrating cells. We integrated T cell subset-specific regulatory annotations from the International Human Epigenome Consortium (IHEC) to infer cellular functions and pathways affected by genomic regions displaying MS-associated differential methylation. Furthermore, to pinpoint expression changes of novel MS-relevant genes across immune cell subtypes, we leveraged locally generated T cell subset datasets. Together, these approaches provide an integrated view of the CSF immune-cell epigenome and reveal MS-associated regulatory programs governing altered immune cell adhesion, migration, and function.

## Results

### CSF cells display typical DNA methylation landscape

To investigate whole-genome DNA methylation changes in people with RRMS, we performed Post-Bisulfite Adaptor Tagging (PBAT)-seq^35^ on CSF cells obtained from age and sex matched RRMS (n=5) and non-inflammatory neurological disease controls (NINDC, n=5), representing the most appropriate and ethically accessible control group for CSF studies (**Fig. 1A and Supplementary Table 1**). To validate the observed methylation differences, we further sequenced a larger cohort including the discovery samples with a lower sequencing depth (**Fig. 1A and Supplementary Table 2**). The discovery samples were utilized to assess concordance between shallow and deep sequencing data, which were treated as technical replicates. The additional independent validation cohort comprising 18 RRMS samples and 12 controls (7 NINDC, and 5 inflammatory neurological disease controls, INDC) was utilized for biological replication. We further used RNA-seq data from overlapping donors^29^ (**Fig. 1A and Supplementary Table 3**), Affymetrix expression data^28^ from an independent larger cohort, IHEC chromatin states annotations and bulk RNA-seq data from sorted T cell subsets to explore the cellular and functional effects (**Fig. 1A**).

**Figure 1.**
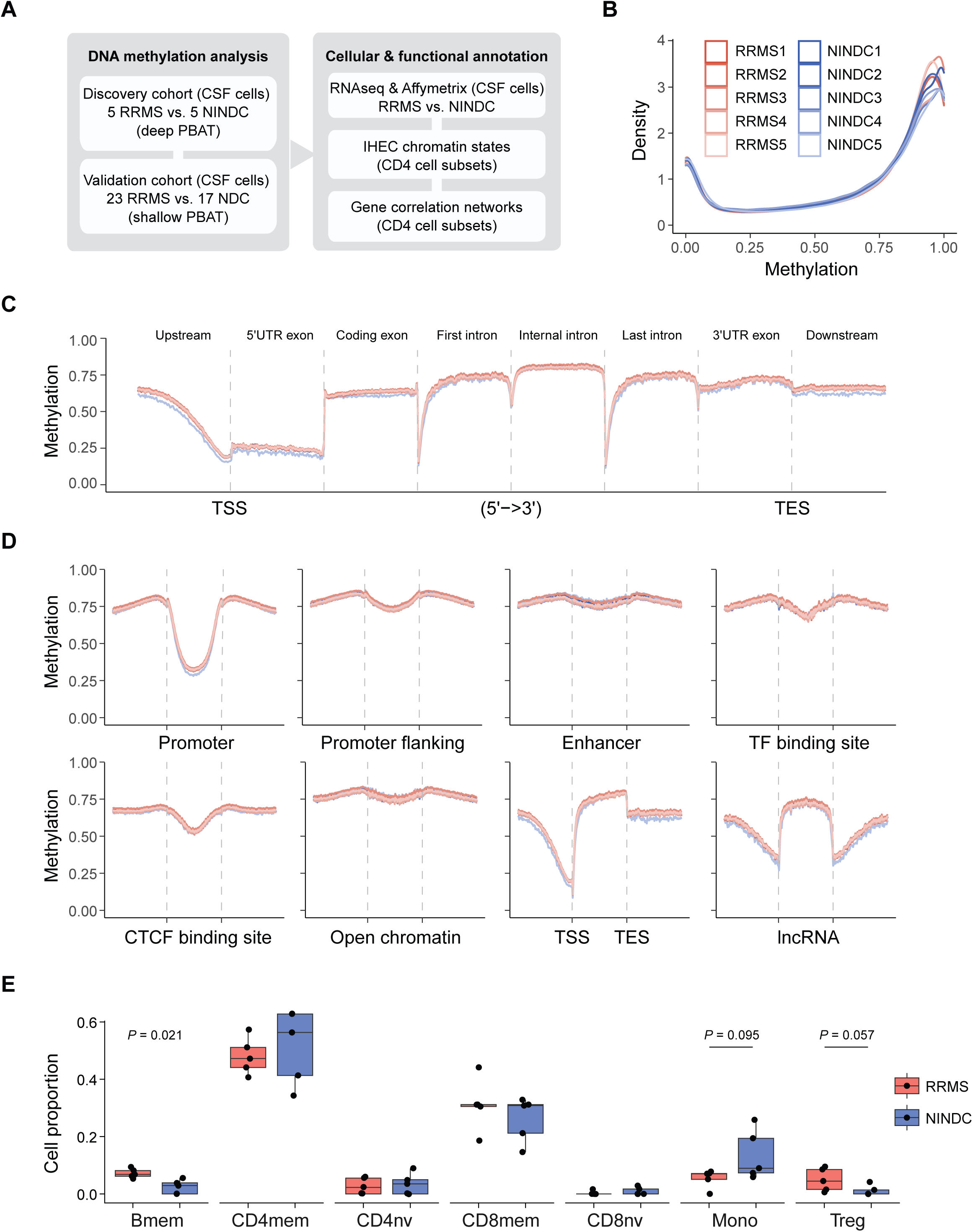
DNA methylation profiling of CSF cells from RRMS and NINDCs. **(A)** Schematic overview of the study design showing CSF cell collection from RRMS and NINDC donors followed by PBAT whole-genome bisulfite sequencing. (B) DNA methylation distribution across RRMS (n=5) and NINDC (n=5) samples (C) DNA methylation patterns across gene-related features. (D) DNA methylation patterns across Ensemble-derived genomic features. The dashed lines indicate the start and end of features. (E) Estimated proportions of immune cell subsets in CSF samples from individuals with RRMS and NINDC. Boxes represent the interquartile range, center lines indicate medians, and whiskers denote 1.5x the interquartile range. Points represent individual samples. Statistical significance between groups was assessed using two-sided Wilcoxon rank-sum tests, with *P* values indicated in each panel. CSF: cerebrospinal fluid, RRMS: Relapsing–remitting multiple sclerosis, NINDC: non-inflammatory neurological disease controls, NDC: neurological disease controls, UTR: untranslated region, TSS: transcriptional start site and TES: transcriptional end site, CTCF: CCCTC-binding factor, lncRNA: long non-coding RNA, Treg: regulatory T cells, CD4Tmem: CD4⁺ memory T cells, CD4Tnv: CD4⁺ naïve T cells, CD8Tmem: CD8⁺ memory T cells, CD8Tnv: CD8⁺ naïve T cells, Bmem: memory B cells, Bnv: naïve B cells, Mono: monocytes.

In the discovering cohort, approximately 28 million CpG sites were sequenced genome-wide (**Supplementary Fig. 1A, B**), and 11.7 million sites remained for analysis after applying a coverage threshold of ≥10x in at least 8 of the 10 samples (**Supplementary Fig. 1C**). Density plots confirmed typical bimodal distribution of methylation with the largest fraction of CpGs being highly methylated (> 75% methylation) and comparably smaller number of CpGs being fully unmethylated (**Fig. 1B**). Low DNA methylation levels (< 25%) were observed within and just upstream of 5’ untranslated regions (UTRs), whereas downstream genic regions displayed high (around 75%) DNA methylation levels (**Fig. 1C**). As expected, promoters and transcription start sites (TSS) showed low methylation, followed by CTCF and transcription factors (TF) binding sites and promoter-flanking regions (**Fig. 1D**). A moderate dip in methylation was also observed in enhancers and open chromatin regions (**Fig. 1D**). Notably, DNA methylation landscape across lncRNA genes resembled that of protein-coding genes (**Fig. 1D**).

Cell-type deconvolution of CSF DNA methylation profiles revealed that T cells constituted the dominant immune population across all samples (**Fig. 1E**). Memory CD4⁺ T cells (CD4Tmem) represented the largest fraction in both groups, followed by CD8⁺ memory T cells. Comparative analysis between RRMS and NINDC samples demonstrated largely comparable overall immune cell compositions. No significant differences were observed for CD4Tmem, CD4Tnv, CD8Tmem or CD8Tnv cells. Monocytes and Tregs showed non-significant trends toward lower and higher levels in RRMS, respectively. A nominally significant difference was detected for memory B cells (Bmem; *P* = 0.021), with higher proportions in RRMS samples. The largest absolute difference in mean cell proportion between groups was 8.08% (monocytes), while the largest median difference was 9.10% (CD4Tmem). Overall, these findings indicate that CSF immune cell composition was broadly comparable between groups, with only modest shifts observed in specific B cell subsets.

Taken together, these results confirm the expected DNA methylation architecture of CSF cells across genomic and regulatory elements, suggest largely comparable immune cell compositions between groups, and support the suitability of PBAT for whole-genome methylome profiling in low-input clinical samples.

### Widespread DNA methylation alterations characterize CSF cells in pwMS

To explore DNA methylation differences between RRMS and NINDC, we applied three complementary analytical methods, Limma, MethylKit, and RADmeth. Each method is built on distinct statistical principles and therefore identified a different number of differentially methylated positions (DMPs), the top significant CpGs were shared (**Supplementary Fig. 2A**). More specifically, the top 1,000 DMPs of each analysis were enriched in the other two methods (**Supplementary Fig. 2A**). In total, 2,710 DMPs were detected with absolute methylation difference ≥ 10% and *P* < 0.001 by minimum two different ways of analysis (**Supplementary Fig. 2B and Supplementary Table 4**). Heatmap and hierarchical clustering of DMPs clearly segregated RRMS from controls, revealing both hyper- and hypomethylated CpGs in RRMS (**Fig. 2A**).

**Figure 2.**
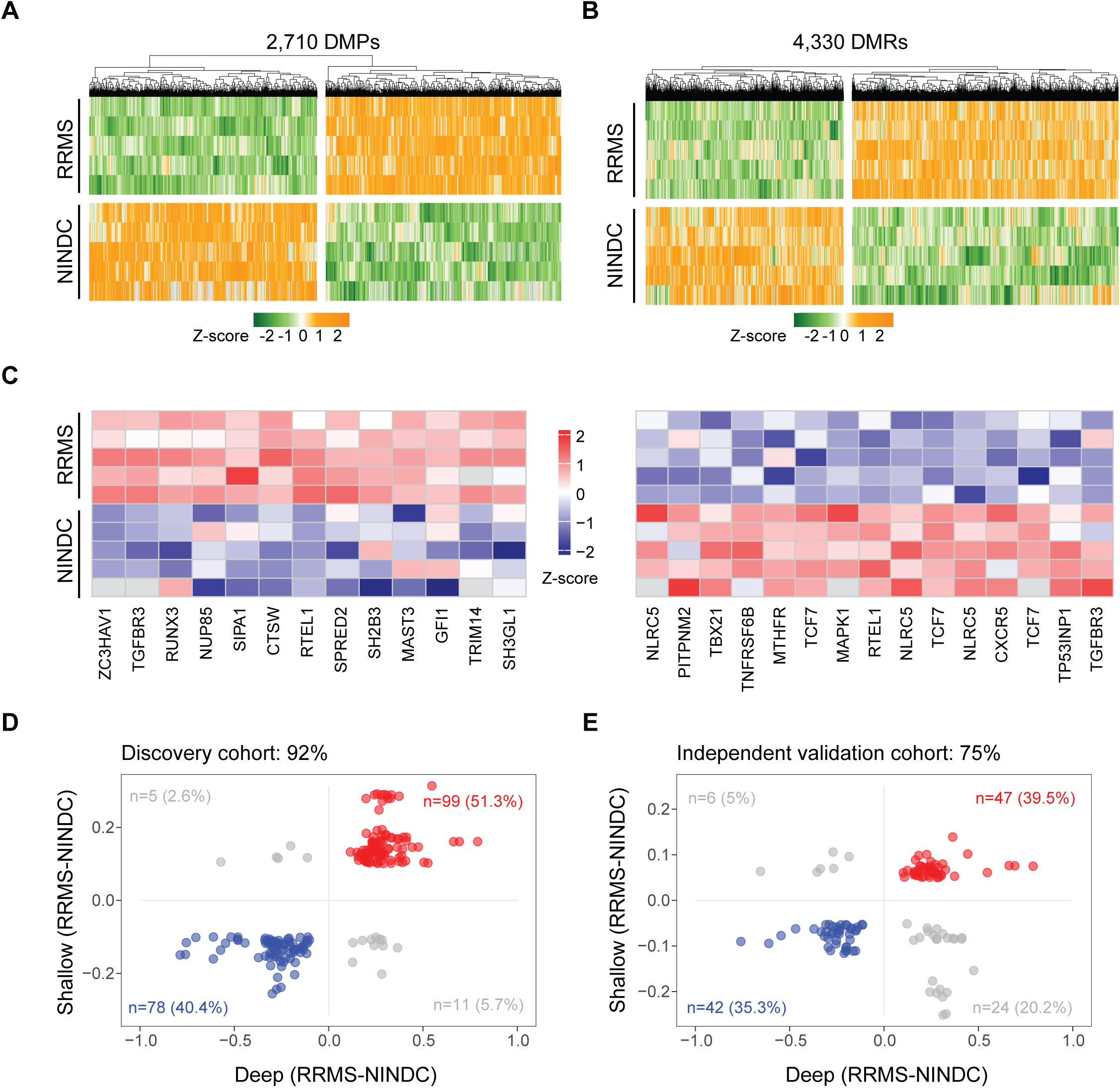
Identification and replication of differential methylation in CSF cells from RRMS and NINDCs. Heatmap and hierarchical clustering of differentially methylated **(A)** positions (DMPs) and **(B)** regions (DMRs). The color-key denotes Z-scored DNA methylation levels (green-to-orange). Replication of DMPs within **(C)** the same (n=10) and **(D)** an independent cohort (n=30) using shallow PBAT-seq. Colors indicate DMPs with the same direction of change in the discovery (deep PBAT-seq) and replication (shallow PBAT-seq) cohorts, with red illustrating increased and blue decreased DNA methylation levels in RRMS. Grey represent DMPs changing in opposite directions. Number of DMPs and (percentages, %) are given in either category in their respective quadrant. DMP: differentially methylated position, DMR: differentially methylated region, RRMS: Relapsing–remitting multiple sclerosis, NINDC: non-inflammatory neurological disease controls.

To increase the robustness of detecting methylation changes with greater functional impact, we next identified differentially methylated regions (DMRs) by clustering CpG sites with a *P* < 0.01 and an absolute methylation difference ≥ 0.1 (see **Methods** and **Supplementary Fig. 2B** for details). In total, 4,330 DMRs were identified (**Fig. 2B and Supplementary Table 5**), with the majority (74%) displaying hypermethylation in RRMS (**Fig. 2B**) and typically spanning < 1kb in length (**Supplementary Fig. 2C**). Methylation levels at the overlapping DMPs and DMRs were highly correlated (ρ > 0.9, **Supplementary Fig. 2D, E**).

To assess the relevance of identified DMPs and DMRs to multiple sclerosis (MS), we overlapped them with reported MS risk genes^5^. DMRs were significantly enriched within 2 kb of MS risk loci (permutation test, *P* = 0.033), with 70 of 560 candidate MS risk genes overlapping DMRs, among which 43 exhibited promoter methylation changes. Gene Ontology analysis of these 70 genes revealed enrichment for pathways related to cytokine responses, regulation of cell adhesion, and T helper (Th) cell differentiation (**Supplementary Table 6**). Further inspection indicated that many of these genes encode chemokines, interleukins, or their receptors, and approximately 30 of the 70 genes have established roles in T cell activation, regulation, and Th differentiation, including processes influencing the balance between Th17 and regulatory T cells (Treg) (**Supplementary Table 6**). After selecting genes with DMRs located in the promoter or flanking regions, we found genes associated with pro-inflammatory Th1/Th17 or activated T cell functions tended to be hypomethylated in MS, including T-box transcription factor 21 (TBX21) and C-X-C chemokine receptor type 5 (CXCR5), whereas genes linked to Treg or anti-inflammatory functions, such as interleukin-2 receptor subunit beta (IL2RB), were more frequently hypermethylated (**Fig. 2C and Supplementary Table 6**). In addition, one DMR (GRCh38: chr19:49,341,982–49,343,506) overlaps a polygenic risk score (PRS)–associated MS risk SNP (chr19:49,343,207) near CD37, a known negative regulator of T cell proliferation³□ (**Supplementary Fig. 3A**).

In the validation cohort, methylation profiles obtained at lower sequencing coverage were smoothed to reduce stochastic noise and improve methylation estimation accuracy^37^ (**Supplementary Fig. 3B, C**). The technical replicates validated 91.7% of testable DMPs, when applying a 10% methylation difference threshold (**Fig. 2D**) and 81.4% with a 5% cutoff (**Supplementary Fig. 3D**). Importantly, biological replication in an expanded independent cohort further confirmed consistent methylation differences for 74.8% of DMPs showing ≥5% methylation change (**Fig. 2E; Supplementary Fig. 3A, B**), a proportion significantly exceeding random expectation (Chi-square test, P = 6.9 × 10⁻□), supporting the robustness of the identified methylation signatures across cohorts.

In summary, integration of results from multiple statistical frameworks and validation in independent samples revealed widespread and reproducible genome-wide DNA methylation alterations in CSF cells from pwMS.

### DNA methylation changes highlight altered T cell adhesion and migration in pwMS

To further characterize MS-associated DMPs and DMRs, we analyzed their distribution and methylation levels across CpG contexts and genomic features. Notably, MS samples exhibited increased methylation in CpG islands, CpG shores, open sea regions, CTCF binding sites, and promoter regions, suggesting a potential repressive effect on gene expression (**Fig. 3A, B, Supplementary Fig. 4A-C**). Notably, while most genomic features showed balanced distributions of hyper- and hypomethylated DMRs, promoter regions displayed a marked enrichment of hypermethylated DMRs in RRMS (**Fig. 3B**). We then used Ingenuity Pathway Analysis (IPA) to explore pathways of genes affected by DMPs and DMRs. The most significant pathways clustered into four categories: G protein-coupled receptors (GPCRs), Rho GTPases/adhesion, Immune/cytokines, and Regulators (**Fig. 3C, Supplementary Table 7, 8**). GPCRs represent a large family of cell surface receptors that transmit extracellular signals into the cell via activation of heterotrimeric G proteins, making them attractive targets for drug development^38^. Interaction of C-X-C motif chemokine receptor type 4 (CXCR4), one of the GPCRs, with its ligand C-X-C motif chemokine ligand 12 (CXCL12) has been reported to guide chemotaxis of T cells and other immune cells, promoting their transmigration across the blood-brain barrier (BBB) and subsequent CNS infiltration in MS^39, 40, 41^. In addition, the GPCR C-C motif chemokine receptor type 7 (CCR7) displayed promoter hypomethylation in pwMS (**Supplementary Table 7**). By primarily mediating immune cell responses to the C-C motif chemokine ligand 19 (CCL19) and C-C motif chemokine ligand 21 (CCL21), it directs naive and memory Th cell entry into lymph nodes for priming and has been implicated in shaping CNS-directed immune responses in MS^42^. Consistently, CCR7 and its ligands are expressed both in peripheral lymphoid tissues and within the CNS in the experimental autoimmune encephalomyelitis (EAE) model^43^. Pathways with ≥10 molecules and BH-adjusted *P* <0.001 were considered in GeneSetCluster2^44^, which identified two main clusters represented by cell adhesion and cell junction assembly (**Supplementary Fig. 4D, Supplementary Table 8**). Furthermore, predicted upstream regulators of DM-associated genes included IL-2, TNF, IFNγ, and TGFB1/TGFBR2, among others (**Fig. 3D, Supplementary Table 9**).

**Figure 3.**
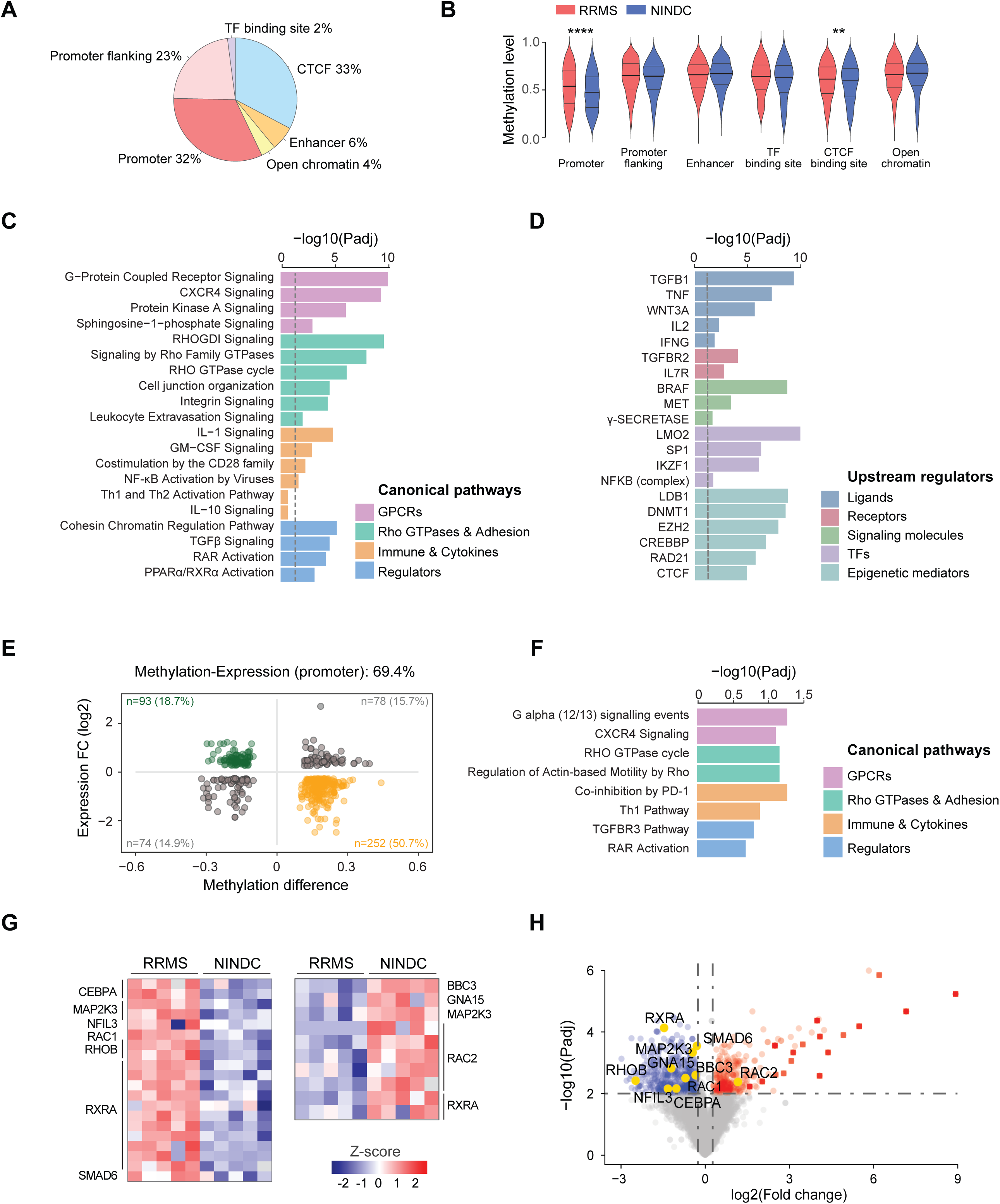
Characteristics of differential methylation and downstream impact on RNA expression. **(A)** Circle plot illustrating the percentage (%) of DMRs across genomic and regulatory features. **(B)** Distribution of DMR methylation levels across genomic and regulatory features in RRMS and NINDC samples. *P* values were calculated using the two-sided Wilcoxon rank-sum tests. The central line indicates the median, and the upper and lower lines represent the 75th and 25th percentiles, respectively. **: *P* < 0.01, ****: *P* < 0.0001 (C-D) Representative enriched pathways **(C)** and upstream regulatory factors **(D)** associated with differentially methylated genes. Bar length indicates enrichment significance (−log10 Padj) and colors represent different canonical pathways. GPCRs: G protein-coupled receptors **(E)** Methylation difference of promoter-associated DMPs and DMRs (RRMS-NINDC) versus the log□ fold changes of corresponding Affymetrix gene expression levels. Genes are categorized according to concordant or inverse methylation–expression changes, in their respective quadrants with numbers (n) and percentages (%) indicated. Green dots indicate genes showing promoter hypomethylation accompanied by increased expression, whereas yellow dots represent genes with promoter hypermethylation and reduced expression. **(F)** Representative canonical pathways enriched among genes exhibiting inverse changes between promoter methylation and gene expression in MS samples. **(G)** Heatmaps of hypermethylated and hypomethylated DMP and DMR-associated genes involved in immune-related and regulatory pathways across RRMS and NINDC samples in **(C)**. **(H)** Volcano plot of log□ fold changes from Affymetrix data, with immune- and regulator-related genes highlighted and labeled. Red and blue dots indicate genes up- or downregulated expression in RRMS (|Foldchange| ≥ 1.2 and Padj < 0.01). Red squares indicate immunoglobulin (IG) genes. DMR: differentially methylated region, DMP: differentially methylated position, RRMS: relapsing–remitting multiple sclerosis, NINDC: non-inflammatory neurological disease controls. Prom: promoter, Prom flk: promoter flanking region, Enh: enhancer, TFbs: transcription factor binding sites, CTCFbs: CTCF binding sites, Open: open chromatin. Dashed line indicates Padj = 0.05 significance threshold.

To investigate the functional consequences of altered methylation in pwMS, we examined transcription using two datasets. We first used RNA-seq of CSF cells from 18 samples (12 RRMS vs 6 NINDC), encompassing 5 and 13 samples used in the discovery (4 RRMS and 1 NINDC) and replication (7 RRMS and 5NINDC) methylation datasets, respectively^29^ (**Supplementary Table 3**). Approximately 69.5% of DMPs and DMRs at promoter regions exhibited inverse methylation and expression changes (**Supplementary Fig. 5A**). More specifically, 47.8% of DMPs and 57.6% of DMRs exhibited hypermethylation in pwMS, accompanied by corresponding decrease in gene expression (**Supplementary Fig. 5A**). We next analyzed Affymetrix microarray data from CSF cells from an independent larger cohort of 26 RRMS and 18 NINDC^28, 45^, and could confirm consistent association between promoter hypermethylation and reduced gene expression in pwMS for nearly half of the altered genes (**Supplementary Fig. 5A**). And the genes with inverse promoter methylation and expression changes were enriched in similar functional categories as the genes associated with DMPs and DMRs (**Fig. 3E, F**). Considering jointly RNA-seq and Affymetrix datasets, 164 genes displayed expression and methylation differences in CSF of pwMS (3.7% of 4,461 genes with DMP or DMR) (**Supplementary Fig. 5B**). Most of these genes displayed hypermethylation and reduced expression. Notably, 12 were related to immune and regulatory pathways, 10 of which overlapped with DEGs identified by both RNA-seq and Affymetrix analyses (**Fig. 3G, H and Supplementary Fig. 5C**). For example, RetinoidLXLReceptorLα (*RXRA*) gene contained multiple hypermethylated sites (**Fig. 3G**) and showed a pronounced reduction in expression in RRMS (**Fig. 3H**). In contrast, *rac family small GTPase 2 (RAC2)* was markedly upregulated in RRMS and exhibited multiple hypomethylated CpGs (**Fig. 3G, H**).

Collectively, these findings indicate that genes involved in T cell adhesion and migration exhibit coordinated changes in both DNA methylation and expression levels in pwMS.

### Differential MS methylation is enriched in active enhancers of memory T cells

To further characterize MS-associated methylation changes, we investigated potential cell type-specific regulation by leveraging cell type-specific chromatin states derived from IHEC. Enrichment analysis across a 100-state universal chromatin model^46^ showed strongest enrichment in the bivalent promoter state 2 (BivProm2), as well as significant enrichment of DMPs and DMRs in EnhA8 and EnhA9, enhancer states typically associated with blood immune cells (**Supplementary Fig. 6A**). Given that CSF cells predominantly consist of T cells, we next examined an 18-state model, which was generated by IHEC using 168 samples comprising diverse T cell subsets. We found the highest enrichment in TSS, particularly upstream flanking (TSSFlnkU), regions repressed by Polycomb (ReprPC) and both active and weak enhancers (EnhA1 and EnhWk) in T cells (**Fig. 4A**). We further conducted enrichment analysis in 229 individual T cell samples from IHEC, demonstrating that while a significant enrichment of MS-associated DMRs was observed in ReprPC across most T cell subsets, more distinct cell type-specific patterns emerged in specific T cell subsets (**Fig. 4B, Supplementary Fig. 6B-C**). Specifically, in naive CD4^+^ T cells, MS-associated DMRs were predominantly enriched in active TSS (TSSA and TSSFlnkU), but in memory CD4^+^ T cell subsets they were mainly enriched in active and weak enhancers (EnhA1, EnhA2 and EnhWk) and downstream flanking TSS (TSSFlnkD) (**Fig. 4B**). Additionally, enrichment in bivalent enhancers (EnhBiv) was the most pronounced in effector and central memory CD4^+^ T cells. In Tregs, the most significant enrichment was in EnhA2, while in Th17 cells, EnhA1, EnhWk and TSSFlnkU, were enriched in MS-associated DMRs. Highly significant enrichment was found in weak enhancers (EnhWk) of immature T cells from thymus and umbilical cord (**Fig. 4B**).

**Figure 4.**
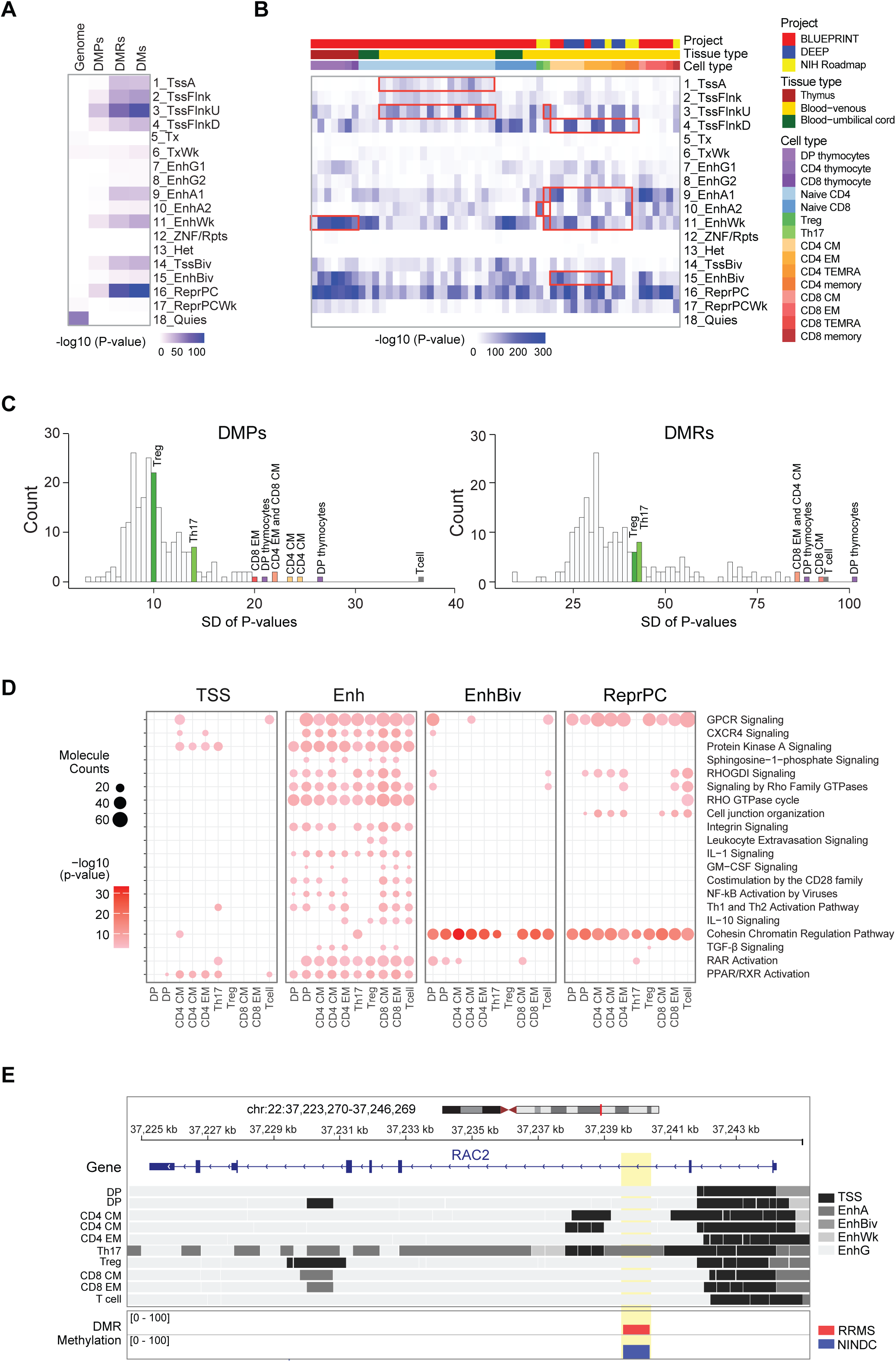
Differential enrichment of DMPs and DMRs across T cell-derived chromatin states. (A-B) Heatmaps showing enrichment of (A) DMPs and DMRs across summary T cell chromatin states and (B) DMRs across selected, representative T cell subtype samples. The color intensity (white-to-blue) represents enrichment significance, −log10(*P*). Top annotation bars indicate project, tissue type, and cell type. Red boxes highlight regions with enrichment differences of interest. (C) Distribution of standard deviation (SD) of enrichment *P* values across all T cell-related chromatin states overlapping with DMPs and DMRs, indicating variability in feature enrichment across T cell subsets. Colored bars and labels denote T cell subtypes with high SD and of particular interest. (D) Representative IPA pathway enrichment across selected T cell subsets within four combined chromatin states: transcription start site (TSS), enhancer (Enh), bivalent enhancer (EnhBiv), and Polycomb-repressed regions (ReprPC). Dot size indicates molecule counts and color intensity (white-to-red) reflects enrichment significance −log10(*P*). Dot color indicates enrichment significance −log10(*P*) and dot size represents molecule counts. (E) A representative genomic view of the RAC2 locus showing localization of DMPs and DMRs (highlighted in yellow) relative to chromatin states across summary and selected T cell subsets. DMP: differentially methylated position, DMR: differentially methylated region, DM: DMPs plus DMRs, 1_TssA: Active transcription start site, 2_TssFlnk: Flanking transcription start site, 3_TssFlnkU: Upstream flanking transcription start site, 4_TssFlnkD: Downstream flanking transcription start site, 5_Tx: Strong transcription, 6_TxWk: Weak transcription, 7_EnhG1: Genic enhancer 1, 8_EnhG2: Genic enhancer 2, 9_EnhA1: Active enhancer 1, 10_EnhA2: Active enhancer 2, 11_EnhWk: Weak enhancer, 12_ZNF/Rpts: ZNF genes and repeats, 13_Het: Heterochromatin, 14_TssBiv: Bivalent/poised transcription start site, 15_EnhBiv: Bivalent enhancer, 16_ReprPC: Polycomb-repressed region, 17_ReprPCWk: Weak Polycomb repression, 18_Quies: Quiescent/low signal region, DP thymocytes: Double-positive thymocytes, CD4 thymocyte: CD4⁺ thymocytes, CD8 thymocyte: CD8⁺ thymocytes, Naive CD4: Naive CD4⁺ T cells, Naive CD8: Naive CD8⁺ T cells, Treg: Regulatory T cells, Th17: T helper 17 cells, CD4CM: CD4⁺ central memory T cells, CD4EM: CD4⁺ effector memory T cells, CD4TEMRA: CD4⁺ terminally differentiated effector memory T cells re-expressing CD45RA, CD4 memory: CD4⁺ memory T cells, CD8CM: CD8⁺ central memory T cells, CD8EM: CD8⁺ effector memory T cells, CD8TEM: CD8⁺ terminally differentiated effector memory T cells re-expressing CD45RA, CD8 memory: CD8⁺ memory T cells.

Based on the distribution of *P* values (**Fig. 4C**), the most heterogeneous enrichment patterns were observed in bulk T cells, double-positive T cells, central and effector memory CD4^+^ and CD8^+^ T cells. To explore potential functional impact of differential methylation in these subsets, including Treg and Th17 (**Fig. 4C, Supplementary Fig. 6D**), we next conducted IPA analysis on the differentially methylated genes mapping to specific chromatin states in a given T cell subset. We found that genes in proximity to MS-associated enhancer-located differentially methylated sites were significantly enriched in pathways including *GPCR signaling*, *Rho GTPases cycle* and *RAR activation* across most T cell subtypes (**Fig. 4D**), suggesting their expression modulation via enhancer methylation in pwMS. A subset of pathways was more specifically associated with promoter (TSS) regions especially in memory CD4^+^ T cells, including *GPCR Signaling*, *PKA Signaling* and *PPAR/RXR activation* (**Fig. 4D**). Interestingly, the *Cohesin Chromatin regulation pathway* demonstrated particularly strong enrichment in EnhBiv and ReprPC regions across all subsets, and active enhancers in Th17 specifically (**Fig. 4D**). While some genes were shared across multiple samples and states, most were uniquely associated with specific cell types and/or chromatin states (**Supplementary Fig. 7**). This is exemplified by the regulatory context of RAC2, where associated DMPs and DMRs reside in genic enhancer (EnhG) regions in bulk T cells and most other cell types, but transition to active enhancer states in Th17 cells, highlighting differentiation state-specific epigenetic regulation (**Fig. 4E**).

Our integrative analysis of MS-associated differential methylation with T cell subset-specific chromatin states suggests a predominant role of enhancer, and to a much lower degree promoter, methylation in regulating T cell activation and migration in pwMS.

### Protocadherin genes are epigenetically repressed in pwMS and implicated in immune regulation

While methylation changes in pwMS were distributed genome-wide, a considerable aggregation of changes was observed in Protocadherin (*PCDH*) genes and clusters, with 6 DMPs and 27 DMRs affecting 43 distinct PCDH genes (**Fig. 5A**, **Supplementary Table 4, 5**). Protocadherin genes constitute a large family of cell-adhesion molecules, categorized into clustered (α, β, γ) and non-clustered genes. They are conventionally expressed in the nervous system to regulate neuronal adhesion, connectivity, and circuit formation^47^, while their role in immune cells remains largely unexplored. We observed extensive methylation alterations across multiple *PCDH* genes, with all DMPs and DMRs except one showing hypermethylation in RRMS compared to controls (**Fig. 5A**). Consistently, transcriptomic data revealed modest but widespread reduction of expression of multiple *PCDH* genes in RRMS, with multiple genes displaying similar downregulation (**Fig. 5A**). Moreover, *PCDH* genes were the main component of the *Cohesin chromatin-regulation pathway* that demonstrated striking enrichment of MS-associated methylation changes across T cell subsets and Th17 cells in particular (**Fig. 4D**).

**Figure 5.**
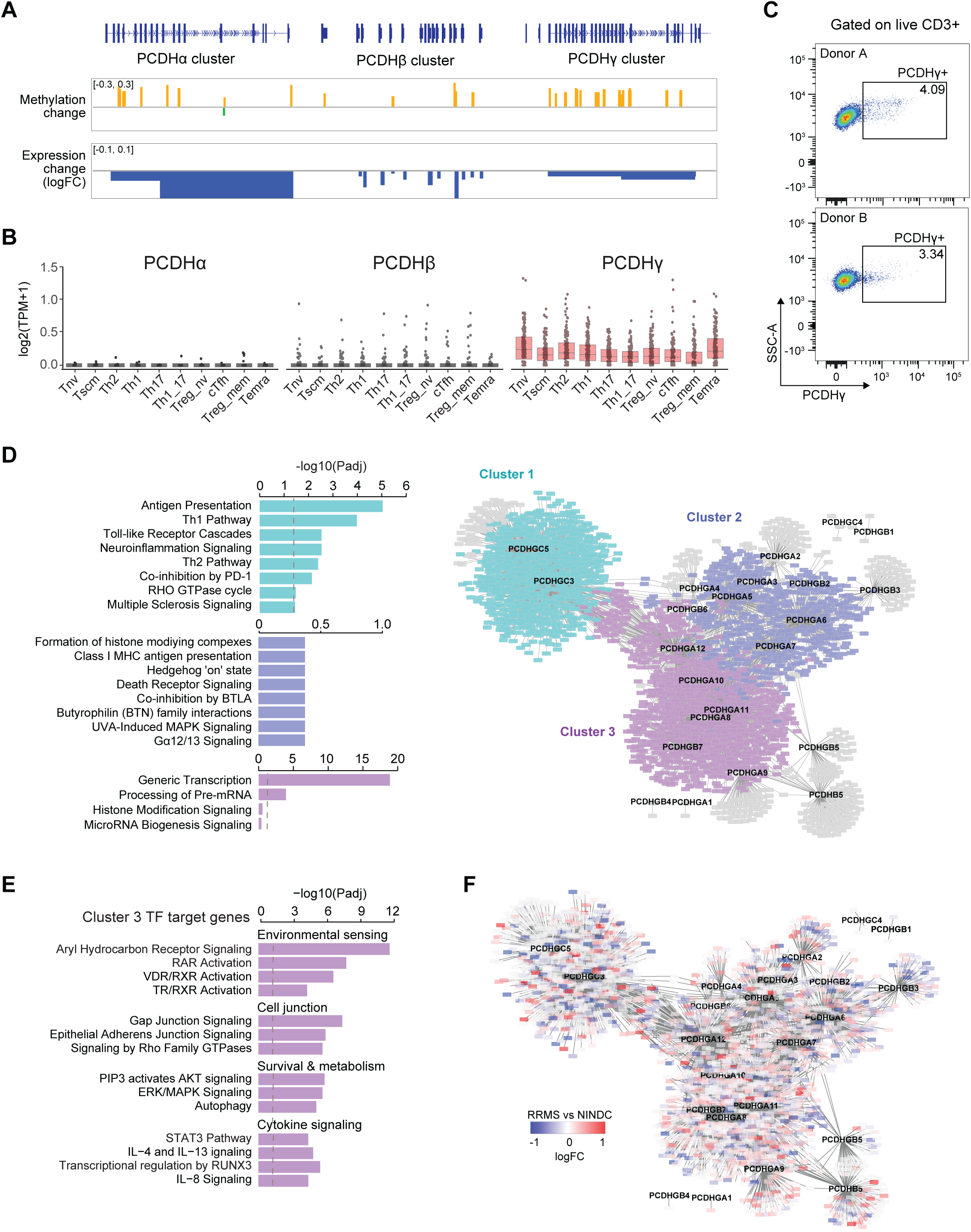
PCDHs are aberrantly methylated and expressed in MS, their co-expression networks and potential functions in T cells. **(A)** DNA methylation differences (RRMS-NINDC) and gene expression changes (log□ fold change) across protocadherin (PCDH) gene clusters. Orange and green bars represent increased and decreased DNA methylation, respectively, and blue bars indicate gene downregulation (logFC). **(B)** Expression patterns of *PCDH*α, *PCDH*β, and *PCDH*γ clusters across ten T cell subsets. Expression values are shown as log□ (TPM+1). Boxplots show median and interquartile range, points indicate individual samples. **(C)** Representative flow cytometry plots showing PCDHγ expression in gated live CD3⁺ T cells from two healthy donors. Cells were analyzed by PCDHγ versus side scatter (SSC-A), with PCDHγ⁺ cells indicated. Percentages represent the proportion of PCDHγ⁺ cells among CD3⁺ T cells. **(D)** Co-expression network of PCDH-associated genes constructed using Spearman correlation. Functional enrichment of major gene clusters is shown on the left. Edges represent significant correlations (|r| ≥ 0.5 and Padj < 0.01). Nodes and pathways are colored according to cluster membership. **(E)** Representative functional categories enriched among transcriptional targets associated with Cluster 3 genes. **(F)** PCDH co-expression network with nodes colored according to gene expression differences (log□ fold change) between RRMS and NINDC samples in Affymetrix expression data. PCDH: protocadherin. FSC-A: forward scatter area, SSC-A: side scatter area. RRMS: relapsing remitting multiple sclerosis NINDC: non-inflammatory neurological disease controls. Dashed line indicates Padj = 0.05 significance threshold.

We next sought to determine whether *PCDH* genes are expressed in T cells, given that they are conventionally associated with neuronal expression and function. To this end, we sorted specific T cell subsets from healthy individuals and performed RNA-seq analysis. Across these subsets, expression of the *PCDH*α and *PCDH*β cluster genes, as well as non-clustered *PCDHs*, was generally low, whereas the *PCDH*γ cluster genes showed appreciable expression (**Fig. 5B**). Notably, *PCDH*γ genes were most expressed in naive T cells (Tnv) and terminally differentiated effector memory T cells re-expressing CD45RA (TEMRA) (**Fig. 5B**). To further validate *PCDH*γ expression at the protein level, we performed intracellular flow cytometry on peripheral blood T cells using an anti-PCDHγ (pan) C-terminal antibody that cross-reacts with *PCDH*γ-A, -B, and -C isoforms. We detected intracellular PCDHγ protein in on average 3-4% of live T cells (**Fig. 5C, Supplementary Fig. 8A, B**), supporting a potential role for *PCDH*γ genes in T cell biology.

To investigate potential functional roles, we performed co-expression analysis between *PCDH* genes and other genes in the T cell subsets. The network genes were segregated into three major clusters. Cluster 1 predominantly comprised first neighbors of *PCDHGC3* and *PCDHGC5*, which showed the highest expression in terminally differentiated effector memory T cells (TEMRA) (**Fig. 5D, Supplementary Fig. 9**). Genes within this cluster were enriched in pathways related to T cell activation and differentiation (**Supplementary Table 10**). Cluster 2 mainly consisted of first neighbors of *PCDHGA5-7* whose function is enriched in activation, chromatin and cell cycle related pathways (**Fig. 5D, Supplementary Fig. 9, Supplementary Table 10**). Cluster 3 was primarily driven by first neighbors of *PCDHGA8,10-11* and *PCDHGB7*, which exhibited relatively lower abundance in Th1_17 cells (**Fig. 5D, Supplementary Fig. 9**). This cluster contained a substantial number of TFs, resulting in the strong enrichment of the transcription and mRNA processing pathways (**Supplementary Table 10**). Examination of TF target genes expressed in T cells further revealed pathways such as *Aryl Hydrocarbon Receptor Signaling*, *RAR Activation*, *VDR*/*RXR Activation*, *mTOR Signaling*, among others (**Fig. 5E, Supplementary Table 10**). Genes co-expressed with PCDH genes in the Affymetrix dataset were enriched for *cAMP-mediated signaling* and *G protein–coupled receptor signaling pathways* (**Supplementary Fig. 8C**).

To further focus on CD4⁺ T cells, the major immune population in CSF, we analyzed single-cell CSF data from RRMS and NINDC samples and projected pseudobulk CD4 gene logFC values onto the co-expression network. Genes co-expressed with PCDHs showed an overall tendency toward reduced expression in RRMS, although this pattern was not uniform across all network components (**Fig. 5F**).

Together, these co-expression analyses identified multiple *PCDH*-centered gene modules that exhibit distinct expression profiles across T cell subsets and are associated with specific transcriptional and signaling programs of relevance in T cells from pwMS.

## Discussion

In this study, we present a comprehensive DNA methylation analysis of CSF cells. Our findings reveal distinct CSF methylation patterns in pwMS with numerous reproducible differentially methylated regions compared to other neurological disease controls. These alterations were predominantly enriched in pathways related to cytokine signaling, immune activation, and cell adhesion and migration, and were accompanied by corresponding changes in RNA expression. Interestingly, abundant methylation changes affected *PCDH* genes and clusters, highlighting a previously unrecognized role of protocadherins in pwMS, and leading to plausible mechanistic connections that may link environmental cues to immune regulation.

We identified numerous methylation changes in regulatory regions of genes strongly enriched in GPCR signaling, Rho GTPase pathways, cytokine signaling, and regulators of immune activation, suggesting enhanced migratory potential and altered communication of CSF immune cells in pwMS. Since we focused on methylation differences larger than 10%, it is unlikely that our findings are driven by bias in cell type composition, as our deconvolution results (**Fig. 1E**), together with single-cell and flow cytometry analyses, indicate that differences in immune cell proportions between MS and controls are substantially smaller than this threshold^32, 33^. Instead, the observed methylation changes likely reflect intrinsic epigenetic alterations within T cells that constitute on average 90% of CSF cells. Our findings, at the epigenetic level of regulation, reinforced the support for enhanced migratory and adhesive properties of immune CSF cells, which may facilitate their ability to cross the BBB and are consistent with the established roles of GPCRs in promoting T cell chemotaxis, BBB transmigration, and CNS infiltration in MS^40, 48^. Importantly, by mapping these alterations across regulatory features, we show that they are not randomly distributed but are preferentially localized to enhancers of genes enriched in these pathways. This enrichment was particularly pronounced in memory CD4^⁺^ T cells, indicating that cells with a high migratory/inflammatory potential are epigenetically primed via specific regulations that are likely causal to corresponding expression changes instead of being a consequence of gene usage. These data uncover a cell type-specific and functionally oriented epigenetic architecture in which DNA methylation at regulatory regions contributes to cytokine signaling, GPCR–Rho GTPase pathways, adhesion, and migration programs in CSF T cells of pwMS. As an example, we identified coordinated DNA methylation and transcriptional alterations in key regulators including *RAC2*, which showed promoter-associated methylation changes accompanied by increased expression in RRMS. *RAC2* encodes a small GTPase with an established role in cytoskeletal remodeling, T cell motility, and autoimmune inflammation^49, 50^. Importantly, a large fraction of genes showed concordant methylation and expression changes, with promoter hypermethylation generally associated with reduced gene expression in pwMS (**Fig. 3E**). Together, these results suggest that CSF pathogenic immune cells in MS are epigenetically programmed toward enhanced migration and adhesion, potentially supporting their accumulation within the CNS inflammatory niche.

In addition to alterations affecting genes and pathways associated with cell trafficking, DNA methylation alterations in CSF immune cells converged on genes and pathways regulating the balance between pathogenic Th1/Th17 responses and regulatory T cell (Treg) programs. Several genes implicated in pro-inflammatory T cell states displayed promoter hypomethylation in pwMS. Those include for example *T-box transcription factor 21 (TBX21)* gene encoding a key transcription factor associated with Th1 differentiation^51^, *adhesion G protein-coupled receptor E5 (ADGRE5)* encoding CD97, which is commonly expressed in activated and inflammatory T cells in MS^52^, and *interleukin-32 (IL32)*, a pro-inflammatory cytokine reported to interact with IL-17 charactering Th17 cells^53^ (**Supplementary Tables 4 and 5**). Consistently, *interleukin-1 receptor accessory protein (IL1RAP)* was also hypomethylated in pwMS, along with significant enrichment of DMPs and DMRs across genes of *IL-1 signaling pathway* (**Supplementary Tables 8 and 9**). IL-1 signaling is a central driver of pathogenic T cell differentiation and CNS trafficking in MS, sustaining inflammatory activation and effector T cell responses^54^. In contrast, several MS candidate genes with established immunoregulatory functions exhibited promoter hypermethylation in CSF cells of pwMS. These included *transforming growth factor beta receptor 3 (TGFBR3)*, which modulates TGF-β signaling and has been implicated in limiting Th17-type inflammatory responses in EAE^55^, and *interleukin-10 receptor subunit alpha (IL10RA*), which mediates *IL-10 signaling*, a central pathway in Treg-mediated immunoregulation^56^ (**Supplementary Tables 7 and 8**). This inflammatory bias may be partially counteracted by alterations in retinoic acid receptor–retinoid X receptor (RAR–RXR) signaling, with *retinoic acid receptor alpha (RARA)* displaying promoter hypermethylation in pwMS (**Supplementary Tables 7 and 8**), which has been reported to promote regulatory T cell differentiation and inhibit Th17 cell development^57^.

An intriguing finding was the widespread hypermethylation across multiple PCDH genes in pwMS, accompanied by coordinated reduction in gene expression in CSF immune cells, as protocadherins are primarily known for their roles in neuronal cell adhesion and their function in immune cells remains poorly understood. Consistent with our observations, independent studies of MS-discordant monozygotic twins have reported altered methylation across PCDH loci, including hypermethylation of the PCDHγ cluster and PCDH10 in PBMCs of affected twins^58^. The *PCDH* gene family of adhesion molecules is critical for neuronal connections and wiring^59^. Various extracellular signals can converge on the same intracellular signaling pathway through the common CD domain shared by members of PCDHα and PCDHγ^60, 61, 62,63^. Mechanistically, isoforms of PCDHα and PCDHγ can be cleaved by γ-secretase and A disintegrin and metalloprotease (ADAM10), subsequently releasing the intracellular domain, which translocate into the nucleus and regulates gene expression^64, 65, 66, 67^. In addition, the intracellular domains of PCDHα and PCDHγ can bind to several kinases involved in cytoskeleton dynamics and cell adhesion, including focal adhesion kinase (FAK), proline-rich tyrosine kinase 2 (Pyk2), and receptor tyrosine kinase rearranged during transformation (Ret)^62, 68, 69, 70, 71^. Together, these findings suggest that protocadherin dysregulation may extend beyond neuronal contexts and contribute to immune-associated regulatory processes. Accordingly, non-clustered PCDHs have been reported to regulate T cell infiltration in cancers^72, 73^ and display altered expression following EBV infection in B cells^74^. In addition, PCDHGA10 is among the targets of primed DNA loops that facilitate rapid gene activation in memory B cells^75^, while PCDHGA1 has been reported as a driver for cell aging^76^. Additional evidence linking protocadherin loci to immune-related processes comes from the identification of methylation changes in PCDHB14 in relation to other inflammatory conditions^77^. However, the expression and functional roles of protocadherins, particularly clustered PCDHs, in immune cells remain largely unexplored warranting future studies.

Supporting a functional role in immune cells, we observed close co-expression between PCDH genes and transcriptional regulators, and flow cytometry confirmed intracellular PCDHγ expression in T cells. Furthermore, γ-secretase emerged as a predicted upstream regulator of differentially methylated genes in our analysis (**Fig. 3D**). Distinct expression patterns of γ-secretase subunits were observed across T cell subsets (**Supplementary Fig. 8D**). In addition, we observed downregulation of one γ-secretase subunit, presenilin 1 (PSEN1, Padj < 0.05, FC ≤ −1.2), in pwMS. These observations collectively suggest altered regulation of γ-secretase-mediated signaling and potential downstream transcriptional effects linked to protocadherins. Interestingly, MS-associated methylation changes were enriched in the *Cohesin Chromatin regulation pathway*, in which *PCDH* genes constituted a substantial component. Given that cohesin is a key mediator of chromatin looping and its anchoring is regulated by CTCF, which functions as a boundary element constraining enhancer–promoter interactions, altered methylation at CTCF motifs may disrupt regulatory domain organization. Interestingly, the enrichment of the cohesin pathway was most pronounced with differentially methylated positions residing in enhancers of Th17 cells, contrasting with preferential localization at bivalent enhancers e in all T cell subsets except Tregs. Together, this suggests that dysregulation of *PCDH* may be linked to epigenetic perturbation of the cohesin-CTCF axis and altered 3D regulatory networks in pro-inflammatory T cells, particularly in Th17 encephalitogenic cells.

These observations jointly support a model in which epigenetic dysregulation of protocadherin loci, altered γ-secretase signaling, and perturbation of higher-order chromatin organization converge to reshape transcriptional programs in T cells in multiple sclerosis. To explore potential downstream mechanisms, we analyzed the pathways targeted by those transcriptional regulators and found strong enrichment in the *Aryl hydrocarbon receptor (AHR) signaling pathway*, with *AHR* displaying hypermethylation and AHR repressor (*AHRR*) exhibiting promoter hypomethylation in pwMS. AHR is a ligand-activated transcription factor residing in the cytoplasm under resting conditions that translocate to the nucleus upon activation and regulate gene expression. AHR ligands are diverse and include xenobiotics present in environmental pollutants, endogenous metabolic products such as tryptophan derivatives, and microbiota- or diet-derived metabolites, making AHR a key player in the integration of environmental and metabolic signals into immune regulation^78^. AHR can direct T cell differentiation toward either Treg or Th17 lineages in response to distinct ligands which can consequently ameliorate or exacerbate MS-like disease in mice^79, 80^. Importantly, consistent with *AHRR* promoter hypomethylation observed in CSF cells of pwMS, smoking, known to predispose both to MS development and to more severe disease, induces pronounced hypomethylation and transcriptional upregulation of the *AHRR* gene in blood immune cells of pwMS^81^. Together, these observations suggest that PCDH-associated transcriptional programs converge on AHR signaling that intersects environmental, metabolic, and viral cues (including EBV activity) and influences immune regulation and potentially contributes to MS pathogenesis.

Despite the strengths of our comprehensive CSF methylome profiling and multi-cohort validation, several considerations remain. Validation across more diverse populations will be essential to confirm the robustness and generalizability of the identified methylation signatures. In addition, although our analyses implicate coordinated *PCDH* silencing by DNA methylation and integration with γ-secretase and AHR signaling, dedicated experimental studies will be required to establish causality and elucidate the underlying mechanisms. Moreover, our analyses were performed on bulk CSF immune populations, which precludes precise attribution of these alterations to specific immune subsets; future single-cell epigenomic and transcriptomic profiling will therefore be important to resolve cellular heterogeneity and define subset-specific contributions. Longitudinal studies, ideally spanning relapse-remission transitions and therapeutic intervention, may further clarify whether these methylation signatures reflect dynamic disease activity or treatment response.

In summary, this study provides the first comprehensive genome-wide characterization of the DNA methylation landscape of CSF immune cells from pwMS, revealing robust and biologically meaningful epigenetic alterations that converge on pathways regulating immune activation, cell migration, and transcriptional control. Our findings highlight enhanced migratory and adhesive programming of CSF immune cells and uncover a previously unknown role of protocadherins and their integration with γ-secretase and AHR signaling, suggesting a mechanistic axis through which environmental, metabolic, and viral cues may shape immune dysregulation in MS. These results not only advance our understanding of MS pathogenesis but also nominate candidate pathways, regulatory networks, and epigenetic signatures with potential relevance for biomarker development, precision patient stratification, and therapeutic targeting.

## Material and Methods

### Cohorts

CSF cells from 5 RRMS and 5 NINDC individuals were included in the genome-wide post-bisulfite adaptor tagging (PBAT) discovery cohort. Technical validation using a less deep WGBS PBAT sequencing was performed on the same samples, as well as on an additional independent cohort 18 RRMS and 7 NINDC and 5 INDC. Of those, 11 RRMS and 6 NINDC CSF cell samples were profiled with RNA sequencing (RNA-seq)^29^. All pwMS patients were diagnosed according to 2010 revised McDonald’s criteria. Patients’ Expanded Disability Status Scale (EDSS) and the Multiple Sclerosis Severity Score (MSSS) were determined by the treating neurologist. Detailed demographic and clinical characteristics of the study participants are provided in Supplementary Table 1-3. The study was approved by the regional ethical committee (ethical permit 2009/2107-31/2) and written informed consent was obtained from all participants.

### Preparation of CSF cells

The CSF was collected in 2 x 15 ml size falcon tubes and centrifuged immediately after lumbar puncture at 440g for 10 minutes at RT to separate the cells, which were subsequently pooled and concentrated to a volume of 20-60 µl per sample. The CSF cells were then frozen immediately on dry-ice and stored in 2 ml size polypropylene tube at -80°C until use.

### Library preparation

DNA methylation libraries were prepared using the PBAT method^35^ with some modifications. Lysates from ∼5000 CSF cells were bisulfite converted using the Imprint DNA Modification Kit (Sigma) by incubation at 99°C for 6 min, 65 °C for 80 min, 95 °C for 3 min and 65 for 20 min. Purification was done according to the manufacturer’s protocol. DNA was eluted in 10 mM Tris-Cl (pH 8.5) and mixed with 0.4 mM dNTPs, 0.4 uM oligo 1 (Biotin)CTACACGACGCTCTTCCGATCTNNNNNNNNN), 1X NEBuffer 2 with a final reaction volume of 24 µl. The samples were incubated at 65 °C for 3 min followed by 4 °C pause, 5 U of Klenow exo- (New England Biolabs) was added and the incubation was continued at 4 °C for 5 min, +1 °C/15 s to 37 °C, 37 °C for 90 min. Samples were then incubated with 20 U exonuclease I (NEB) for 1 h at 37 °C. DNA was purified with 0.8X Agencourt Ampure XP beads (Beckman Coulter), eluted in 10 mM Tris-Cl (pH 8.5) and incubated with washed M-280 Streptavidin Dynabeads (Invitrogen) for 30 min with rotation at room temperature. Beads were washed twice with 0.1 N NaOH, and twice with 10 mM Tris-Cl (pH 8.5) and resuspended in 47 µl of 0.4 mM dNTPs, 0.4 µM oligo 2 (TGCTGAACCGCTCTTCCGATCTNNNNNNNNN) and 1X NEBuffer 2. Samples were then incubated at 95 °C for 45 s followed by 4 °C pause before addition of 10 U Klenow exo- and then incubated at +1 °C/15 s to 37 °C, 37 °C for 90 min. Beads were washed with 10 mM Tris-Cl (pH 8.5) and resuspended in 50 µl of 0.4 mM dNTPs, 0.4 µM PE1.0 forward primer (AATGATACGGCGACCACCGAGATCTACACTCTTTCCCTACACGACGCTCTTCCGATCT), 0.4 µM indexed iPCRTag reverse primer, 1 U KAPA HiFi HotStart DNA polymerase (KAPA Biosystems) in 1X HiFi Fidelity Buffer. Libraries were amplified by PCR as follows: 95 °C 2 min, 12-15 cycles of (98 °C 80 s, 65 °C 30 s, 72 °C 30 s), 72 °C 3 min and 4 °C hold. Amplified libraries were purified using 0.8X Agencourt Ampure XP beads. Libraries’ quality and quantity were determined using High-Sensitivity DNA chips on the Agilent Bioanalyzer, and the KAPA Library Quantification Kit for Illumina (KAPA Biosystems). Pools of three libraries were prepared for HiSeq 100 bp Single-end sequencing on a HiSeq2500 in High Output mode (8 lanes/run) at the Babraham Institute Next Generation Sequencing Facility. 5 RRMS and 5 NINDC samples were sequenced with 150 bp paired end reads on 20 HiSeqX lanes (2 lanes per sample) at the National Genomics Infrastructure (NGI, Stockholm).

### Trimming and filtering of sequencing reads

Single end reads of 100 bp were processed using Trim Galore! (v0.4.1) developed by Babraham Bioinformatics, with the first 9 bp clipped from the 5′ end (--clip_R1 9). Quality (-q 20) and adapter (-a AGATCGGAAGAGC) trimming were performed using Cutadapt (v1.8.1)^82^ with a minimum required adapter overlap of 1bp (-O 1) and a maximum trimming error rate of 0.1 (-e 0.1). Reads shorter than 20 bp after trimming were discarded. Trimmed and filtered sequencing reads were aligned to GRCh38 with the -pbat ^35^ option of Bismark (v0.14.4)^83^, using Bowtie 2 and the following specified options: -q -phred33 -score-min L,0,-0.2 -ignore-quals. Alignments with a unique best hit were taken for further processing.

### Estimation of DNA methylation

Duplicated reads were removed using the deduplicate_bismark function and subsequently top and bottom strands were merged into single CpG dinucleotides using the -merge_CpG option of the Bismark coverage2cytosine module^83^. Methylation level values were calculated by reads mapped as cytosine divided by total coverage. CpG with coverage >=10 in more than 8 samples were kept for downstream analysis and methylation level were NA if the coverage is less than 10. Chromosome X and Y were removed. Gene and genomic feature annotations were obtained from Ensembl^84^, GRCh38, gene were extended upstream for 2kb. BEDTools^85^ were used to intersect the CpGs. Methylation pattern analyses were performed by CpGtools^86^.

### Cell-type proportion estimation

Immune cell-type composition in cerebrospinal fluid (CSF) samples was estimated by reference-based deconvolution of DNA methylation data using EpiDISH (v2.22.0; CBS method)^87^. We used the cent12CT.m reference matrix comprising 12 immune cell subtypes^88^. Reference CpG probes were lifted from hg19 to hg38 coordinates and matched to CpG sites covered in each sample. Methylation beta values overlapping reference CpGs were retained for deconvolution, resulting in variable CpG overlap across donors (112–513 CpGs). Cell-type fractions were estimated using the epidish function and reported as proportions of the 12 immune cell subtypes. Estimated proportions between RRMS and NINDC were compared using two-sided Wilcoxon rank-sum tests. Cell-type proportions are visualized as boxplots showing medians, interquartile ranges, and individual samples.

### Identification of DMPs

Limma regression, which uses empirical Bayes moderation, allowing us to consider co-variants such as age and sex^89^. MethylKit utilizes chi-square test^90^, while RADmeth takes advantage of Beta-binomial distribution^91^. We utilized all three methods to identity DMP candidates. For Limma and RADmeth, the previously mentioned filtering strategy, which retained CpG sites with a minimum coverage of 10X in at least eight samples, was used. In the case of methylKit, a uniform minimum coverage threshold of 10X was applied across all samples, as the method requires consistent coverage. Sex and age were included as covariates in both Limma and methylKit models. CpG sites which met the criteria of *P* < 0.001 and absolute methylation differences ≥ 0.1 in at least two of the three methods were defined as DMPs.

### Identification of DMRs

To identify DMRs, CpG sites with a *P* < 0.01 and an absolute methylation difference ≥ 0.1 determined by any of the three methods, were selected as candidate sites for DMRs. Candidate CpGs were then clustered by BEDTools with a maximum inter-CpG distance of 1,000 bp, applying a strand-specific mode (the direction of methylation level change). Regions containing more than three candidate CpG sites were retained and defined as DMRs.

### Enrichment analysis of DMRs near MS risk loci

MS risk loci were defined based on lead SNPs reported by the International Multiple Sclerosis Genetics Consortium^5^. Genomic coordinates were harmonized to the hg38 reference genome using UCSC liftOver. Each SNP was extended to a 4kb window centered on the lead variant (±2 kb). Enrichment of DMRs overlapping these regions was assessed using a permutation-based approach. DMR coordinates were randomly shifted 5,000 times across the genome while preserving region length, and the number of overlaps was calculated for each iteration. An empirical *P* value was derived as the proportion of permutations yielding overlap counts greater than or equal to the observed value.

### Replication cohort data analysis

To improve precision and enable the use of CpG sites with lower coverage, weighted local regression (BSmooth)^37^ was applied with default parameters to smooth methylation profiles, with weights determined by read coverage. Smoothed methylation levels were used as input to estimate differential methylation between RRMS and INDC and NINDC control samples. CpG sites exhibiting consistent directional changes across both deep and shallow sequencing datasets, for the 10 technical replicate samples, were used as input for the independent cohort comparison, where quadrant plots were generated.

### DMP and DMR’s annotation

Gene and genomic feature annotations were obtained from Ensembl^84^, based on the GRCh38 reference genome. Gene coordinates were extended 2kb upstream of the transcriptional start site to include promoters. CpG sites were intersected with genomic features using BEDtools^85^. CpG island annotations were downloaded from UCSC Genome Browser hg38^92^. DNA methylation patterns across genomic features were explored using CpGtools^86^. EpiLogos was used to visualize epigenomic feature around DMRs overlapping MS risk SNPs^93^.

### Function analysis and regulator prediction

Genes associated with DMPs and DMRs were subjected to Ingenuity Pathway Analysis (IPA, QIAGEN) for functional annotation and upstream regulator prediction^94^. Enrichment significance was determined using Benjamini-Hochberg (BH)-adjusted *P* values. Significantly enriched pathways (Padj < 0.05 and molecule count ≥ 10) were grouped using GeneSetCluster (version 2.0)^44^. IPA was also applied to explore subsets of genes with 1) differential promoter methylation and inverse expression changes, 2) DMP and DMR associated genes within different IHEC features and cell types, 3) genes within PCDH clusters, and 4) target genes of generic transcriptional regulatory pathways. Genomic tracks were visualized using the Integrative Genomics Viewer^95^.

### IHEC enrichment and functions

Individual chromatin states from 229 T cell samples (healthy donors), as well as summary chromHMM annotations were extracted from IHEC. Enrichment analysis was performed using the ChromHMM OverlapEnrichment (-center) tool^96^ with default parameters. UpsetR^97^ was used to visualize overlapping genes across different T cell types and genomic features We next conducted IPA analysis on the differentially methylated genes mapping to four chromatin state groups TSS, Enh, EnhBiv and ReprPC in a given T cell subset. The final example of RAC2 was shown in integrative genomics viewer (IGV)^95^.

### Flow cytometric staining of pan-PCDH**γ**

Human peripheral blood mononuclear cells (PBMCs) were stained using a pan-PCDHγ antibody panel and analyzed on a BD LSRFortessa flow cytometer. Briefly, 1 × 10L cells were incubated with a fixable viability dye and Fc receptor block. Cells were then fixed with 3.65% formaldehyde for 20 min and permeabilized with 0.1% IGEPAL CA-630 for 4 min to enable intracellular detection of pan-PCDHγ. A mouse anti-pan-PCDHγ antibody (abcam, cat.no. ab187186), was used to detect the C-terminus region common to multiple members of the PCDHγ family, and signal was amplified by staining with a secondary AF488-conjugated anti-mouse IgG antibody (abcam, cat.no. ab150113). Subsequently, staining of lineage-defining surface markers was carried out using antibodies against CD3 (BioLegend, cat.no. 300328), CD20 (Miltenyi, cat.no. 130-111-338).

### PCDH co-expression network

Transcript per million (TPM) values from RNA-seq data of naïve conventional CD4⁺ T cells (Tnv), T stem cell memory T cells (Tscm), T helper 2 cells (Th2), T helper 1 cells (Th1), T helper 17 cells (Th17), T helper 1/17 cells (Th1_17), circulating T follicular helper cells (cTfh), naïve regulatory T cells (Treg_nv), memory regulatory T cells (Treg_mem), terminally differentiated effector memory T cells re-expressing CD45RA (Temra) were used to generate the network. Genes with a summed TPM ≥ 1 across all samples were retained for correlation analysis. Pairwise gene correlations were calculated using Spearman’s rank correlation coefficient. Gene pairs with an absolute correlation coefficient ≥ 0.5 and a Padj < 0.01 were used to construct the co-expression network. Network visualization and construction were performed using Cytoscape^98^. Transcription factor–target regulons were obtained from the DoRothEA human database (dorothea_hs), using interactions with confidence levels A–C^99^.

Robust Multi-array Average (RMA) normalization was applied to Affymetrix CSF microarray data to generate expression values for supplementary network construction^28, 45^. Genes with a summed expression value ≥ 10 were retained for downstream analysis. Pairwise gene correlations were calculated using Pearson correlation, and associations with the Padj < 0.001 were used for IPA. Pathways with an IPA enrichment Padj < 0.0001 were included in the network.

## Data availability

The methylation sequencing data in this study is available in European Genome-phenome Archive (EGA) database.

## Acknowledgements

The authors acknowledge support from the National Genomics Infrastructure in Stockholm funded by Science for Life Laboratory, the Knut and Alice Wallenberg Foundation and the Swedish Research Council, and SNIC/Uppsala Multidisciplinary Center for Advanced Computational Science for assistance with massively parallel sequencing and access to the UPPMAX computational infrastructure. Processing of sequencing data including quality control, filtering, and read alignment, was enabled by resources provided by the National Academic Infrastructure for Supercomputing in Sweden (NAISS) at UPPMAX, funded by the Swedish Research Council through grant agreement no. 2022-06725. The authors thank the Babraham Institute Genomics Facility and the Babraham Institute Bioinformatics Unit for assistance with Illumina sequencing and data analysis. We thank the International Human Epigenome Consortium (IHEC) for providing access to harmonized and reprocessed epigenomic data spanning a diverse range of human cell and tissue types.

This study was supported by grants from the Swedish Research Council, the Swedish Association for Persons with Neurological Disabilities, the Swedish Brain Foundation, the Swedish MS Foundation, the Stockholm County Council (ALF project), StratNeuro, Neuroförbundet, the European Research Council grant (grant agreement No 818170) and the Knut and Alice Wallenberg Foundation. M. Jagodic has received funding from the EU Horizon2020, European Research Council, Swedish Research Council, Swedish Brain Foundation. L.K. is supported by a fellowship from the Margaretha af Ugglas Foundation. Work in GK’s lab was funded by the Biotechnology and Biological Sciences Research Council (BBS/E/B/000C0423) and Medical Research Council (MR/K011332/1). The Babraham Bioinformatics Unit and Genomics Facility were supported by the Babraham Institute’s BBSRC Core Capability Grant (BB/CCG2210/1).

## Author contributions

Y.H. performed the majority of data analyses and drafted the manuscript. G.Y.Z. and M.P.K. prepared the PBAT sequencing libraries. C.S. designed and performed the flow cytometry experiments, H.L. and Mi.K. performed the flow cytometry data analysis. N.R. and C.R.P. generated the T cell subset datasets, and V.B. generated the corresponding data matrices.

E.I. and G.K. assisted with the PBAT sequencing libraries. N.H. performed the single cell analyses. Mo.K., F.P., and T.O. contributed to patient recruitment, sample collection, and clinical data interpretation. L.K. contributed to data analysis and manuscript revision. M.N. performed the shallow-sequencing analyses, revised the manuscript, and co-supervised the project. M.J. conceptualized and designed the study, secured funding, supervised the project, and revised the manuscript. All authors read and approved the final manuscript.

## Competing interests

The authors declare no competing interests.

---

**Supplementary Figure 1. Overview of samples and sequencing depth.**

**(A)** Total number of CpG sites covered per sample at 10X sequencing depth. **(B)** Number of retained CpG sites across samples under different minimum coverage thresholds. **(C)** Number of retained CpG sites as a function of both minimum CpG coverage and minimum sample inclusion thresholds.

**Supplementary Figure 2. Comparison of differential methylation analysis methods.**

**(A)** Enrichment analysis of the top 1,000 DMPs identified by Limma (first column), methylKit (second column), RADmeth (third column), and the final DMP set (fourth column), evaluated across ranked *P* values from Limma (first row), methylKit (second row), and RADmeth (third row). **(B)** Overlap of DMP candidates detected by Limma, methylKit, and RADmeth (*P* < 0.001; mean absolute methylation difference ≥ 0.1). **(C)** Distribution of DMR lengths. **(D)** Overlap and correlation between DMPs and DMRs. **(E)** Correlation of methylation levels between DMPs and DMRs. DMP: differentially methylated position, DMR: differentially methylated region.

**Supplementary Figure 3. Coverage of shallow-sequenced samples and replication of DMPs.**

**(A)** Heatmap showing the number of retained CpG sites across samples at varying coverage thresholds. **(B)** Line plot illustrating CpG site retention across ten sample groups at increasing minimal coverage thresholds. **(C)** Visualization of the DMR, MS-risk SNPs, and IHEC chromatin-state annotations at the CD37 locus. Chromatin state colors follow the standard ChromHMM 18-state annotation. **(D)** Quadrant distribution of CpG methylation differences between shallow and deep sequencing. Axes indicate methylation differences between datasets, and the shallow WGBS were filtered by an absolute methylation difference threshold of 0.05. Numbers indicate counts and percentages of CpGs in each quadrant.

**Supplementary Figure 4. Characteristics and functional annotation of differentially methylated positions (DMPs) and regions (DMRs).**

**(A)** Proportional distribution of DMPs across CpG features and genomic annotations (left), and methylation levels of DMPs within these features in RRMS and NINDC samples (right). *P* values were calculated using the two-sided Wilcoxon rank-sum test. **(B)** Proportional distribution (top) and methylation levels (bottom) of DMRs across CpG features. *P* values were calculated using the two-sided Wilcoxon rank-sum test, with asterisks indicating statistical significance. The central line in (A-B) indicates the median, and the upper and lower lines represent the 75th and 25th percentiles, respectively. **(C)** Proportional distribution of background CpG sites across CpG features (top) and genomic annotations (bottom). **(D)** GeneSetCluster analysis of pathways enriched among genes associated with DMRs (adjusted *P* < 0.05 and gene count ≥ 10). Heatmap shows similarity among enriched pathways with hierarchical clustering. ****: *P* <1 × 10⁻^4^. DMP: differentially methylated position, DMR: differentially methylated region.

**Supplementary Figure 5. Overlap between DNA methylation and gene expression changes.**

**(A)** Scatter plots showing DNA methylation differences plotted against gene expression fold changes (log2) in RNA-sequencing and Affymetrix datasets. Analyses were performed separately for promoter-associated and all gene-associated DMPs and DMRs. Points are colored by quadrant, representing concordant or discordant directions of methylation and expression changes; numbers and percentages indicate the proportion of features in each quadrant. **(B)** Overlap between differentially methylated genes and DEGs. **(C)** Overlap between differentially methylated genes and DEGs within inflammation-related pathways. DMP: differentially methylated position, DMR: differentially methylated region.

**Supplementary Figure 6. Enrichment of DMPs and DMRs across distinct IHEC chromatin states.**

**(A)** Enrichment of DMPs and DMRs across 108 IHEC chromatin states. Color intensity represents enrichment significance −log10(*P*). **(B)** Enrichment of DMPs and DMRs across all available IHEC T cell samples, with chromatin states shown on rows and samples on columns. Top **(C)** Enrichment of DMPs across IHEC T cell samples, showing a representative subset of naïve CD4⁺ T cell samples for visualization clarity. **(D)** Enrichment of DMPs and DMRs across ten representative IHEC T cell samples selected based on the highest variability in chromatin-state enrichment across samples, together with Th17 and Treg samples. Annotation bars indicate project, tissue type, and cell type for (B-D). DMP: differentially methylated position, DMR: differentially methylated region, DM: DMPs plus DMRs, 1_TssA: Active transcription start site, 2_TssFlnk: Flanking transcription start site, 3_TssFlnkU: Upstream flanking transcription start site, 4_TssFlnkD: Downstream flanking transcription start site, 5_Tx: Strong transcription, 6_TxWk: Weak transcription, 7_EnhG1: Genic enhancer 1, 8_EnhG2: Genic enhancer 2, 9_EnhA1: Active enhancer 1, 10_EnhA2: Active enhancer 2, 11_EnhWk: Weak enhancer, 12_ZNF/Rpts: ZNF genes and repeats, 13_Het: Heterochromatin, 14_TssBiv: Bivalent/poised transcription start site, 15_EnhBiv: Bivalent enhancer, 16_ReprPC: Polycomb-repressed region, 17_ReprPCWk: Weak Polycomb repression, 18_Quies: Quiescent/low signal region, DP thymocytes: Double-positive thymocytes, CD4 thymocyte: CD4⁺ thymocytes, CD8 thymocyte: CD8⁺ thymocytes, Naive CD4: Naive CD4⁺ T cells, Naive CD8: Naive CD8⁺ T cells, Treg: Regulatory T cells, Th17: T helper 17 cells, CD4CM: CD4⁺ central memory T cells, CD4EM: CD4⁺ effector memory T cells, CD4TEM: CD4⁺ effector memory T cells re-expressing CD45RA, CD4 memory: CD4⁺ memory T cells, CD8CM: CD8⁺ central memory T cells, CD8EM: CD8⁺ effector memory T cells, CD8TEM: CD8⁺ effector memory T cells re-expressing CD45RA, CD8 memory: CD8⁺ memory T cells.

**Supplementary Figure 7. Gene overlaps across ten T cell samples within four IHEC genomic features.**

UpSet plots showing overlaps of genes associated with differential methylation across ten T cell samples within four IHEC genomic features: TSS, enhancer (Enh), bivalent enhancer (EnhBiv), and repressive Polycomb (ReprPC). Bar plots indicate the size of gene intersections, while the dot matrix denotes the gene numbers of T cell samples contributing to each intersection.

**Supplementary Figure 8. Flow cytometric analysis of PCDHγ expression, co-expression network analysis, and γ-secretase gene expression across T cell subtypes.**

**(A)** Representative gating strategy for flow cytometric analysis of pan-protocadherin gamma (PCDHγ) expression. Cells were sequentially gated on lymphocytes (forward scatter area, FSC-A, vs side scatter area, SSC-A), single cells, live cells, and CD3⁺ T cells, followed by analysis of pan-PCDHγ expression plotted against CD20. **(B)** Flow cytometric analysis of pan-PCDHγ staining under surface-only conditions and after surface staining followed by secondary antibody incubation in CD3⁺ T cells from two donors. Gates indicate pan-PCDHγ⁺ cells, and percentages represent the proportion of pan-PCDHγ⁺ cells among CD3⁺ T cells. **(C)** Boxplots showing RNA expression levels of γ-secretase complex genes (APH1A, APH1B, NCSTN, PSEN1, PSEN2, and PSENEN) across T cell subtypes. Boxplots show the median (center line) and interquartile range (box), whiskers extend to 1.5 × IQR, and points represent individual samples. **(D)** Enriched pathways of PCDH co-expressed genes. Pairwise gene correlations were calculated using Pearson correlation, and significantly co-expressed gene pairs (adjusted P < 0.001) were used for pathway enrichment analysis. **(E)** Co-expression network constructed from genes belonging to significantly enriched Ingenuity Pathway Analysis (IPA) pathways. FSC-A: forward scatter area, SSC-A: side scatter area, PCDHγ: protocadherin gamma, Tconv: conventional T cells, Tscm: T stem cell memory, Tfh: T follicular helper cells, Treg: regulatory T cells, TEMRA: effector memory T cells re-expressing CD45RA, IPA: Ingenuity Pathway Analysis.

**Supplementary Figure 9. Expression of PCDH genes across T cell subsets.**

Supplementary Figure 9. Expression of protocadherin (PCDH) genes across T cell subsets. Boxplots showing log□(TPM + 1) transformed RNA-seq expression levels of expressed PCDH genes with detectable expression across ten T cell subtypes. Boxplots show the median (center line) and interquartile range (box), whiskers extend to 1.5 × IQR, and points represent individual samples.

## Notes

### Competing Interest Statement

The authors have declared no competing interest.

