## Supplementary figures and tables for "Epigenomic profiling of cerebrospinal fluid cells identifies immune regulatory alterations and implicates protocadherins in multiple sclerosis": Sup.Figures.pdf

**Sup. Figure 1**

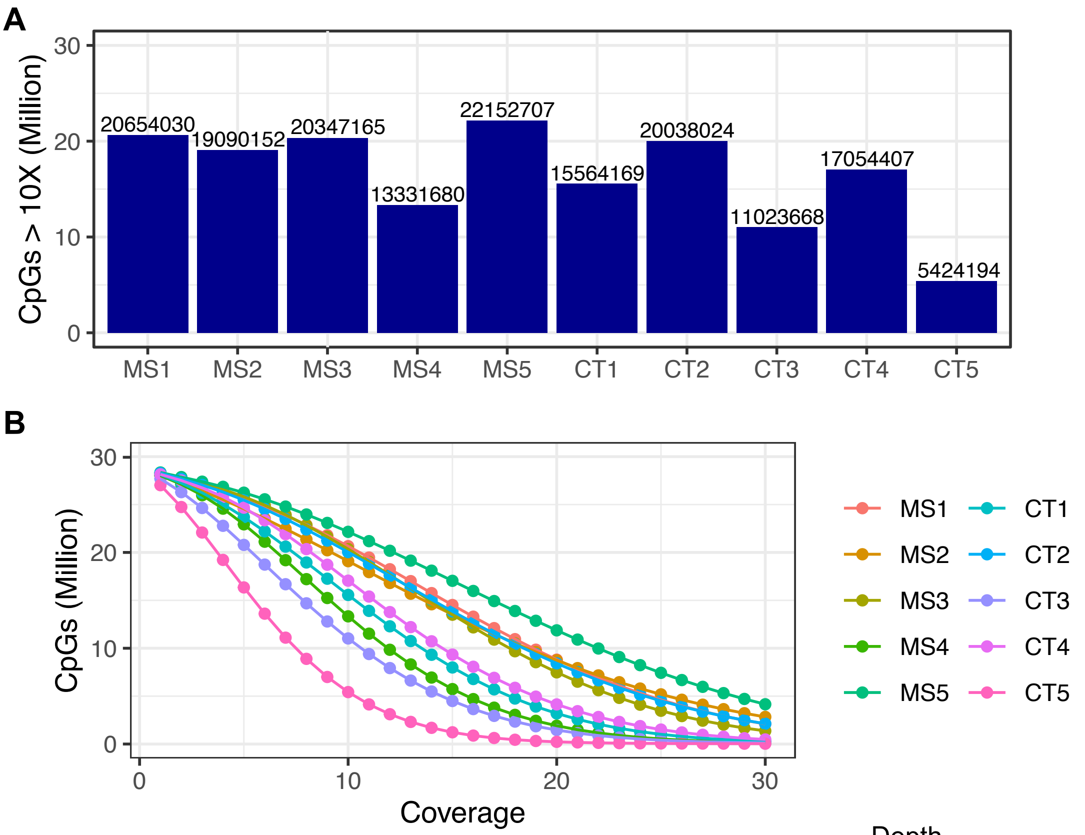


C

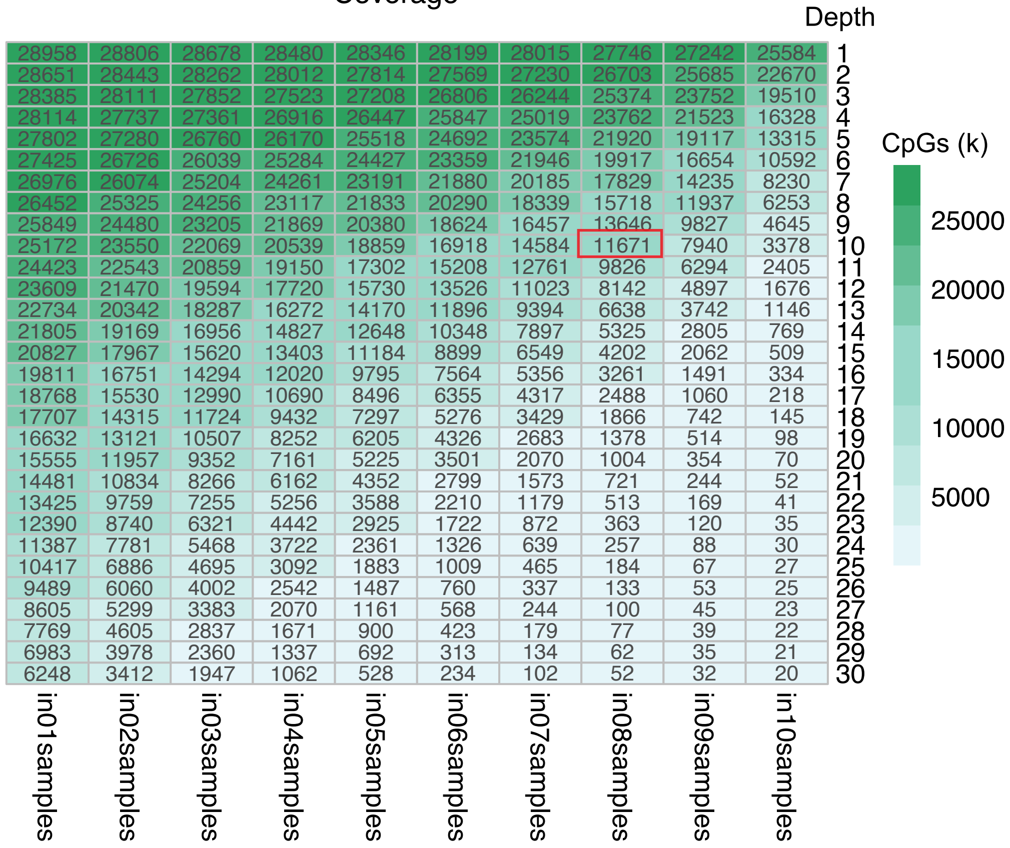


| Depth | in01samples | in02samples | in03samples | in04samples | in05samples | in06samples | in07samples | in08samples | in09samples | in10samples |
|---|---|---|---|---|---|---|---|---|---|---|
| 1 | 28958 | 28806 | 28678 | 28480 | 28346 | 28199 | 28015 | 27746 | 27242 | 25584 |
| 2 | 28651 | 28443 | 28262 | 28012 | 27814 | 27569 | 27230 | 26703 | 25685 | 22670 |
| 3 | 28385 | 28111 | 27852 | 27523 | 27208 | 26806 | 26244 | 25374 | 23752 | 19510 |
| 4 | 28114 | 27737 | 27361 | 26916 | 26447 | 25847 | 25019 | 23762 | 21523 | 16328 |
| 5 | 27802 | 27280 | 26760 | 26170 | 25518 | 24692 | 23574 | 21920 | 19117 | 13315 |
| 6 | 27425 | 26726 | 26039 | 25284 | 24427 | 23359 | 21946 | 19917 | 16654 | 10592 |
| 7 | 26976 | 26074 | 25204 | 24261 | 23191 | 21880 | 20185 | 17829 | 14235 | 8230 |
| 8 | 26452 | 25325 | 24256 | 23117 | 21833 | 20290 | 18339 | 15718 | 11937 | 6253 |
| 9 | 25849 | 24480 | 23205 | 21869 | 20380 | 18624 | 16457 | 13646 | 9827 | 4645 |
| 10 | 25172 | 23550 | 22069 | 20539 | 18859 | 16918 | 14584 | 11671 | 7940 | 3378 |
| 11 | 24423 | 22543 | 20859 | 19150 | 17302 | 15208 | 12761 | 9826 | 6294 | 2405 |
| 12 | 23609 | 21470 | 19594 | 17720 | 15730 | 13526 | 11023 | 8142 | 4897 | 1676 |
| 13 | 22734 | 20342 | 18287 | 16272 | 14170 | 11896 | 9394 | 6638 | 3742 | 1146 |
| 14 | 21805 | 19169 | 16956 | 14827 | 12648 | 10348 | 7897 | 5325 | 2805 | 769 |
| 15 | 20827 | 17967 | 15620 | 13403 | 11184 | 8899 | 6549 | 4202 | 2062 | 509 |
| 16 | 19811 | 16751 | 14294 | 12020 | 9795 | 7564 | 5356 | 3261 | 1491 | 334 |
| 17 | 18768 | 15530 | 12990 | 10690 | 8496 | 6355 | 4317 | 2488 | 1060 | 218 |
| 18 | 17707 | 14315 | 11724 | 9432 | 7297 | 5276 | 3429 | 1866 | 742 | 145 |
| 19 | 16632 | 13121 | 10507 | 8252 | 6205 | 4326 | 2683 | 1378 | 514 | 98 |
| 20 | 15555 | 11957 | 9352 | 7161 | 5225 | 3501 | 2070 | 1004 | 354 | 70 |
| 21 | 14481 | 10834 | 8266 | 6162 | 4352 | 2799 | 1573 | 721 | 244 | 52 |
| 22 | 13425 | 9759 | 7255 | 5256 | 3588 | 2210 | 1179 | 513 | 169 | 41 |
| 23 | 12390 | 8740 | 6321 | 4442 | 2925 | 1722 | 872 | 363 | 120 | 35 |
| 24 | 11387 | 7781 | 5468 | 3722 | 2361 | 1326 | 639 | 257 | 88 | 30 |
| 25 | 10417 | 6886 | 4695 | 3092 | 1883 | 1009 | 465 | 184 | 67 | 27 |
| 26 | 9489 | 6060 | 4002 | 2542 | 1487 | 760 | 337 | 133 | 53 | 25 |
| 27 | 8605 | 5299 | 3383 | 2070 | 1161 | 568 | 244 | 100 | 45 | 23 |
| 28 | 7769 | 4605 | 2837 | 1671 | 900 | 423 | 179 | 77 | 39 | 22 |
| 29 | 6983 | 3978 | 2360 | 1337 | 692 | 313 | 134 | 62 | 35 | 21 |
| 30 | 6248 | 3412 | 1947 | 1062 | 528 | 234 | 102 | 52 | 32 | 20 |

**Sup. Figure 2**

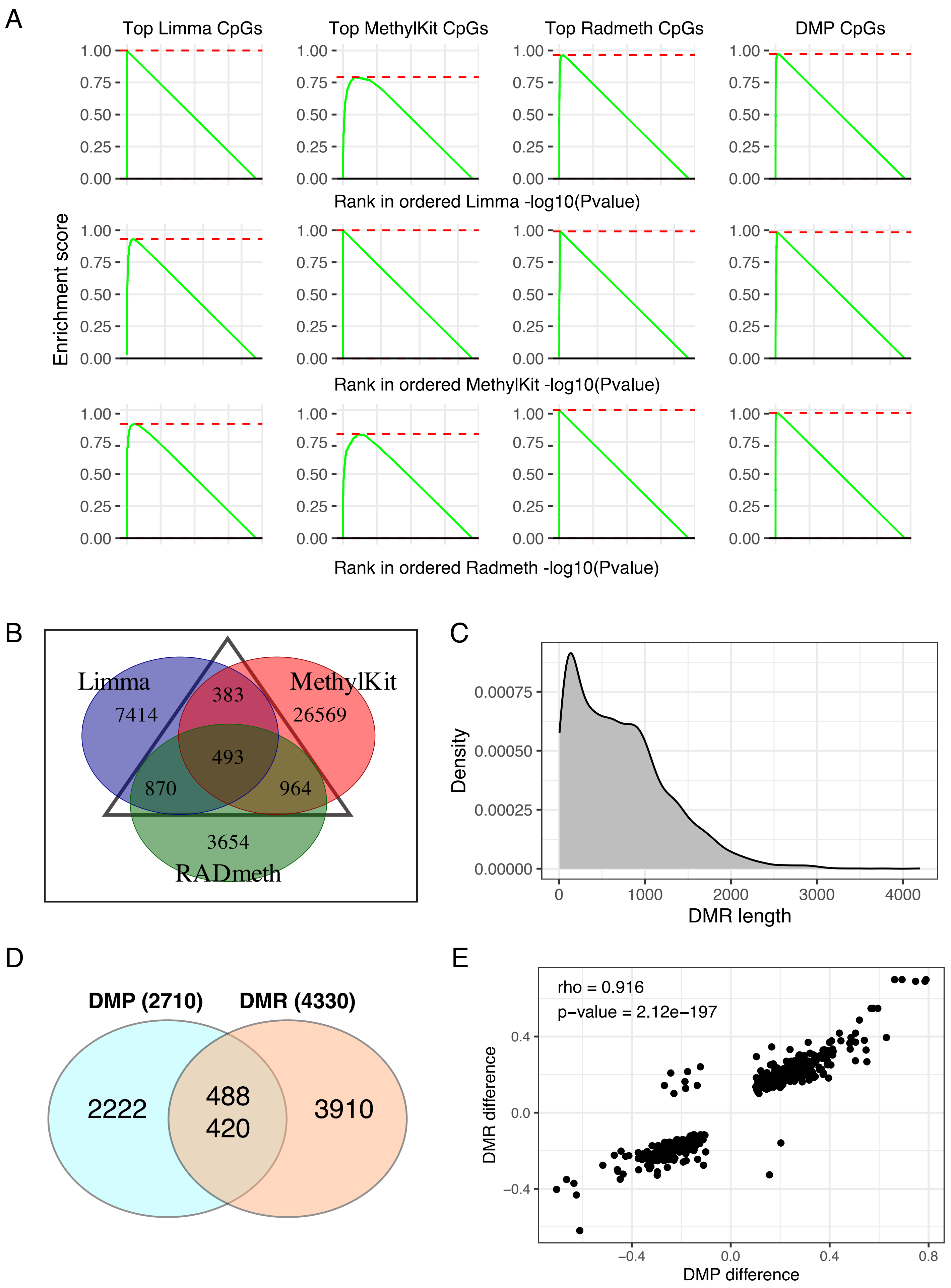
A
Top Limma CpGs
Top MethylKit CpGs
Top Radmeth CpGs
DMP CpGs
Enrichment score
1.00
0.75
0.50
0.25
0.00
Rank in ordered Limma -log10(Pvalue)
Rank in ordered MethylKit -log10(Pvalue)
Rank in ordered Radmeth -log10(Pvalue)
B
Limma
7414
MethylKit
26569
383
493
870
964
3654
RADmeth
C
Density
0.00075
0.00050
0.00025
0.00000
0
1000
2000
3000
4000
DMR length
D
DMP (2710)
DMR (4330)
2222
488
420
3910
E
rho = 0.916
p-value = 2.12e−197
DMR difference
0.4
0.0
−0.4
−0.4
0.0
0.4
0.8
DMP difference

### Sup. Figure 3

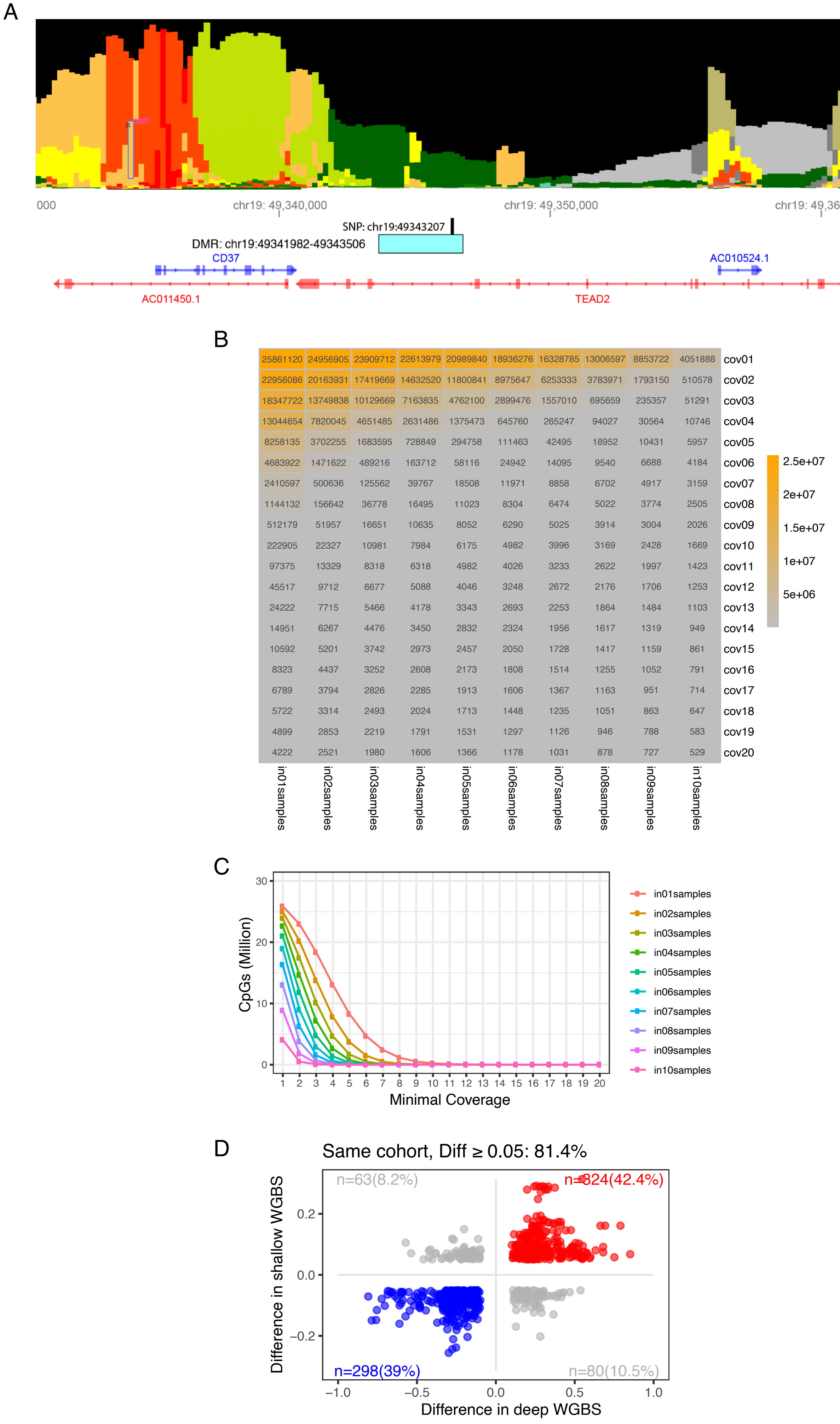
A
000
chr19: 49,340,000
chr19: 49,350,000
chr19: 49,360,000
SNP: chr19:49343207
DMR: chr19:49341982-49343506
CD37
AC010524.1
AC011450.1
TEAD2
B
25861120 24956905 23909712 22613979 20989840 18936276 16328785 13006597 8853722 4051888 cov01
22956086 20163931 17419669 14632520 11800841 8975647 6253333 3783971 1793150 510578 cov02
18347722 13749838 10129669 7163835 4762100 2899476 1557010 695659 235357 51291 cov03
13044654 7820045 4651485 2631486 1375473 645760 265247 94027 30564 10746 cov04
8258135 3702255 1683595 728849 294758 111463 42495 18952 10431 5957 cov05
4683922 1471622 489216 163712 58116 24942 14095 9540 6688 4184 cov06
2410597 500636 125562 39767 18508 11971 8858 6702 4917 3159 cov07
1144132 156642 36778 16495 11023 8304 6474 5022 3774 2505 cov08
512179 51957 16651 10635 8052 6290 5025 3914 3004 2026 cov09
222905 22327 10981 7984 6175 4982 3996 3169 2428 1669 cov10
97375 13329 8318 6318 4982 4026 3233 2622 1997 1423 cov11
45517 9712 6677 5088 4046 3248 2672 2176 1706 1253 cov12
24222 7715 5466 4178 3343 2693 2253 1864 1484 1103 cov13
14951 6267 4476 3450 2832 2324 1956 1617 1319 949 cov14
10592 5201 3742 2973 2457 2050 1728 1417 1159 861 cov15
8323 4437 3252 2608 2173 1808 1514 1255 1052 791 cov16
6789 3794 2826 2285 1913 1606 1367 1163 951 714 cov17
5722 3314 2493 2024 1713 1448 1235 1051 863 647 cov18
4899 2853 2219 1791 1531 1297 1126 946 788 583 cov19
4222 2521 1980 1606 1366 1178 1031 878 727 529 cov20
in01samples in02samples in03samples in04samples in05samples in06samples in07samples in08samples in09samples in10samples
2.5e+07
2e+07
1.5e+07
1e+07
5e+06
C
CpGs (Million)
30
20
10
0
1 2 3 4 5 6 7 8 9 10 11 12 13 14 15 16 17 18 19 20
Minimal Coverage
in01samples
in02samples
in03samples
in04samples
in05samples
in06samples
in07samples
in08samples
in09samples
in10samples
D
Same cohort, Diff ≥ 0.05: 81.4%
n=63(8.2%)
n=324(42.4%)
n=298(39%)
n=80(10.5%)
Difference in shallow WGBS
0.2
0.0
−0.2
−1.0
−0.5
0.0
0.5
1.0
Difference in deep WGBS

**Sup. Figure 4**

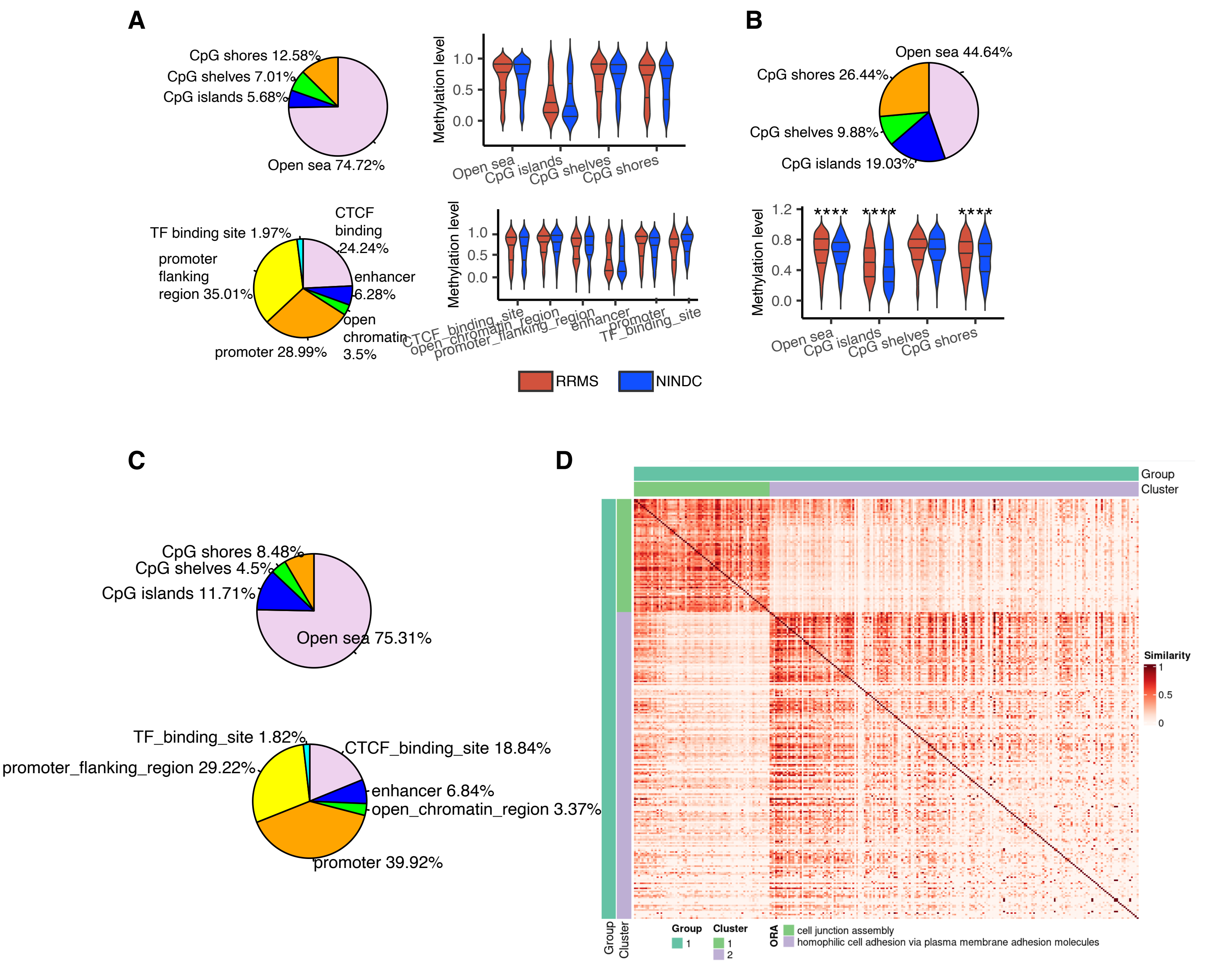
A
CpG shores 12.58%
CpG shelves 7.01%
CpG islands 5.68%
Open sea 74.72%
Methylation level
1.0
0.5
0.0
Open sea
CpG islands
CpG shelves
CpG shores
TF binding site 1.97%
CTCF binding 24.24%
promoter flanking region 35.01%
enhancer 6.28%
open chromatin 3.5%
promoter 28.99%
CTCF_binding_site
open_chromatin_region
promoter_flanking_region
enhancer
promoter
TF_binding_site
RRMS
NINDC
B
Open sea 44.64%
CpG shores 26.44%
CpG shelves 9.88%
CpG islands 19.03%
1.2
0.8
0.4
0.0
****
****
****
C
CpG shores 8.48%
CpG shelves 4.5%
CpG islands 11.71%
Open sea 75.31%
TF_binding_site 1.82%
CTCF_binding_site 18.84%
promoter_flanking_region 29.22%
enhancer 6.84%
open_chromatin_region 3.37%
promoter 39.92%
D
Group
Cluster
Similarity
1
0.5
0
Group
1
Cluster
1
2
ORA
cell junction assembly
homophilic cell adhesion via plasma membrane adhesion molecules

**Sup. Figure 5**

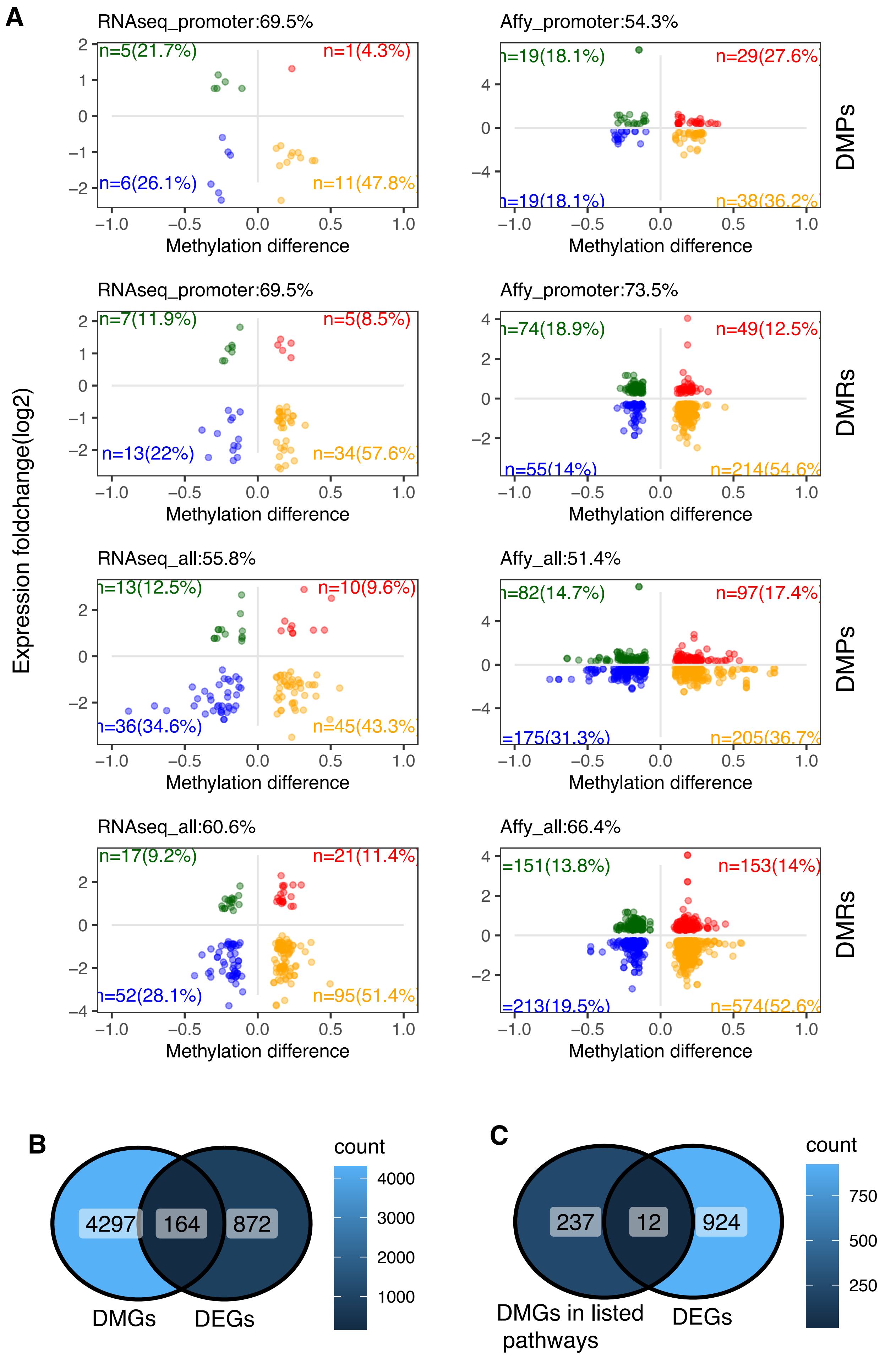
A
RNAseq_promoter:69.5%
n=5(21.7%)
n=1(4.3%)
n=6(26.1%)
n=11(47.8%)
Affy_promoter:54.3%
n=19(18.1%)
n=29(27.6%)
n=19(18.1%)
n=38(36.2%)
DMPs
RNAseq_promoter:69.5%
n=7(11.9%)
n=5(8.5%)
n=13(22%)
n=34(57.6%)
Affy_promoter:73.5%
n=74(18.9%)
n=49(12.5%)
n=55(14%)
n=214(54.6%
DMRs
RNAseq_all:55.8%
n=13(12.5%)
n=10(9.6%)
n=36(34.6%)
n=45(43.3%)
Affy_all:51.4%
n=82(14.7%)
n=97(17.4%)
=175(31.3%)
n=205(36.7%
DMPs
RNAseq_all:60.6%
n=17(9.2%)
n=21(11.4%)
n=52(28.1%)
n=95(51.4%)
Affy_all:66.4%
=151(13.8%)
n=153(14%)
=213(19.5%)
n=574(52.6%
DMRs
Expression foldchange(log2)
Methylation difference
B
4297
164
872
DMGs
DEGs
count
4000
3000
2000
1000
C
237
12
924
DMGs in listed pathways
DEGs
count
750
500
250

#### Sup. Figure 6

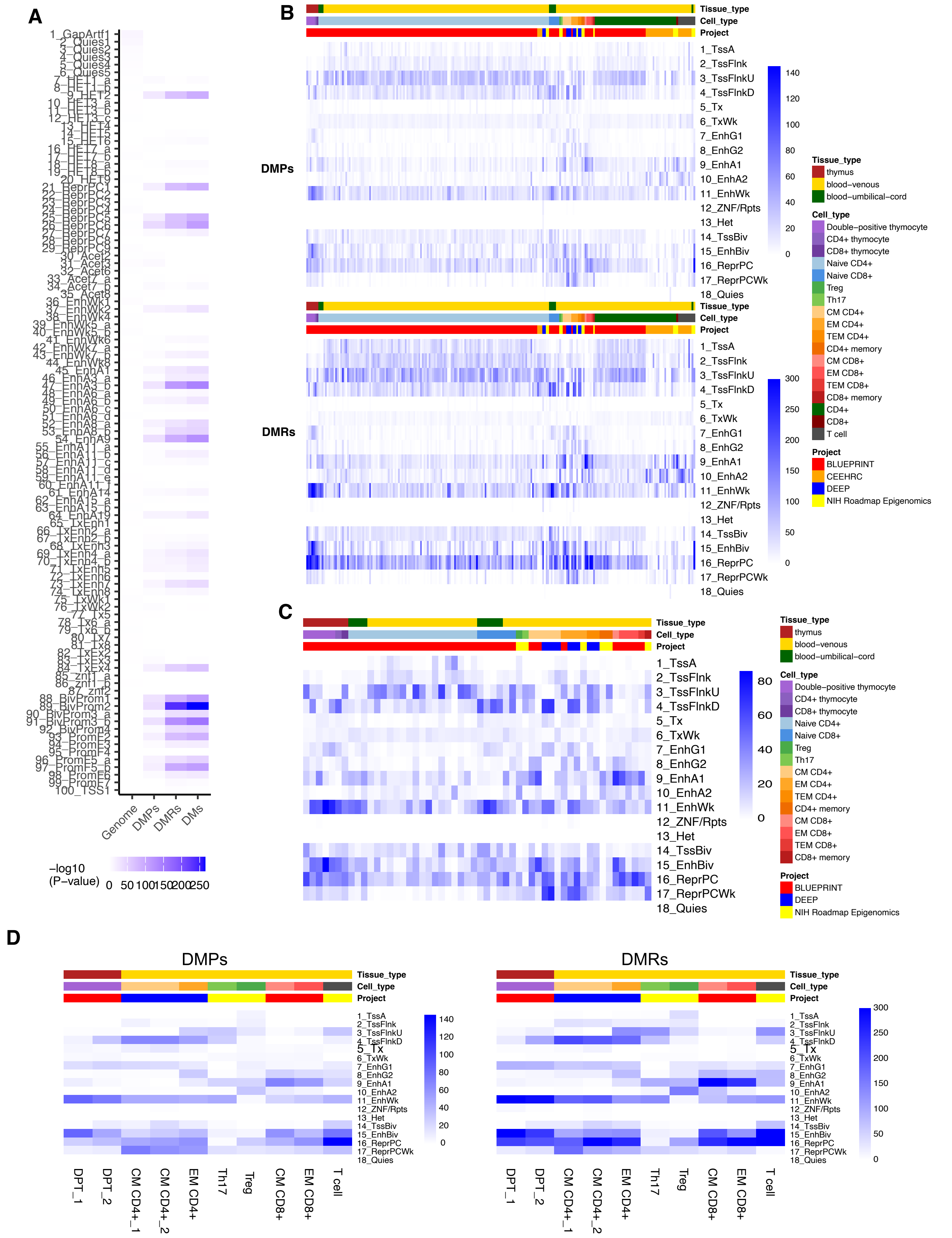
A
B
C
D
DMPs
DMRs
Genome
DMPs
DMRs
DMs
-log10 (P-value)
Tissue_type
Cell_type
Project
thymus
blood-venous
blood-umbilical-cord
Double-positive thymocyte
CD4+ thymocyte
CD8+ thymocyte
Naive CD4+
Naive CD8+
Treg
Th17
CM CD4+
EM CD4+
TEM CD4+
CD4+ memory
CM CD8+
EM CD8+
TEM CD8+
CD8+ memory
CD4+
CD8+
T cell
BLUEPRINT
CEEHRC
DEEP
NIH Roadmap Epigenomics
1_TssA
2_TssFlnk
3_TssFlnkU
4_TssFlnkD
5_Tx
6_TxWk
7_EnhG1
8_EnhG2
9_EnhA1
10_EnhA2
11_EnhWk
12_ZNF/Rpts
13_Het
14_TssBiv
15_EnhBiv
16_ReprPC
17_ReprPCWk
18_Quies
DPT_1
DPT_2
CM CD4+_1
CM CD4+_2
EM CD4+
Th17
Treg
CM CD8+
EM CD8+
T cell

#### Sup. Figure 7

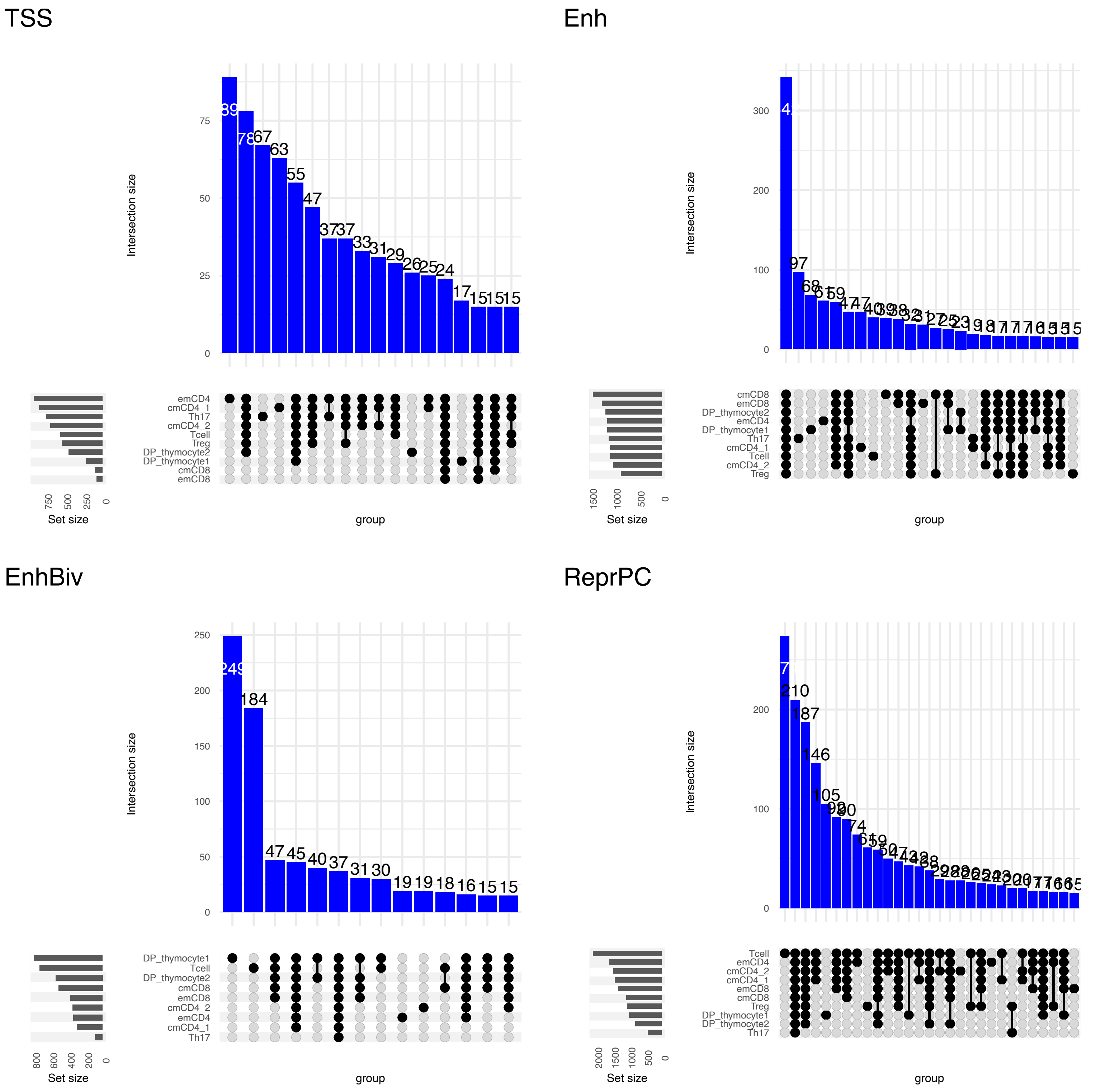

TSS
Intersection size
0
25
50
75
89
78
67
63
55
47
37
37
33
31
29
26
25
24
17
15
15
15
emCD4
cmCD4_1
Th17
cmCD4_2
Tcell
Treg
DP_thymocyte2
DP_thymocyte1
cmCD8
emCD8
750
500
250
0
Set size
group
Enh
Intersection size
0
100
200
300
97
68
cmCD8
emCD8
DP_thymocyte2
emCD4
DP_thymocyte1
Th17
cmCD4_1
Tcell
cmCD4_2
Treg
1500
1000
500
0
Set size
group
EnhBiv
Intersection size
0
50
100
150
200
250
249
184
47
45
40
37
31
30
19
19
18
16
15
15
DP_thymocyte1
Tcell
DP_thymocyte2
cmCD8
emCD8
cmCD4_2
emCD4
cmCD4_1
Th17
800
600
400
200
0
Set size
group
ReprPC
Intersection size
0
100
200
210
187
146
105
90
74
Tcell
emCD4
cmCD4_2
cmCD4_1
emCD8
cmCD8
Treg
DP_thymocyte1
DP_thymocyte2
Th17
2000
1500
1000
500
0
Set size
group

**Sup. Figure 8**

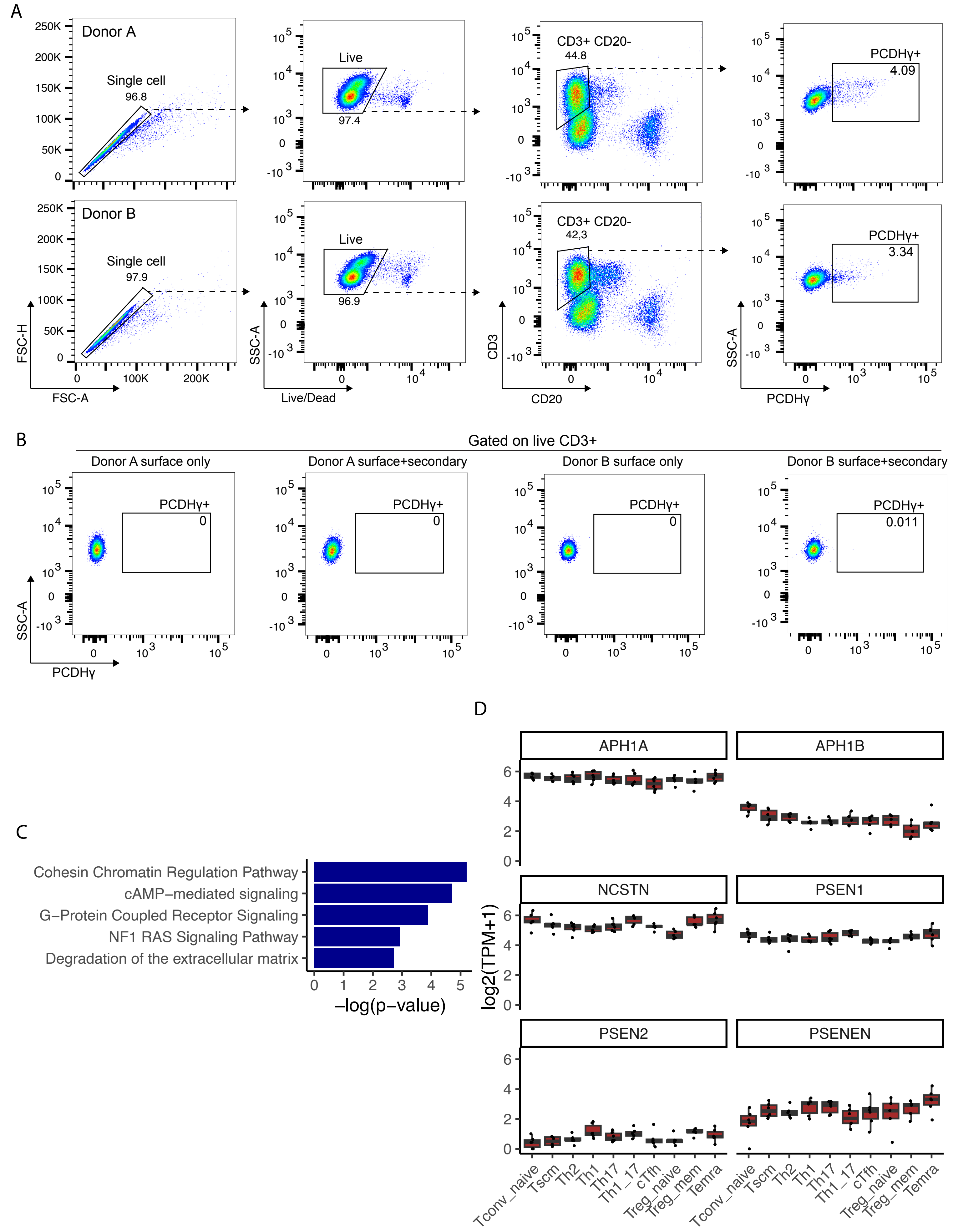
A
Donor A
Single cell
96.8
Live
97.4
CD3+ CD20-
44.8
PCDHγ+
4.09
Donor B
Single cell
97.9
Live
96.9
CD3+ CD20-
42,3
PCDHγ+
3.34
FSC-H
FSC-A
SSC-A
Live/Dead
CD3
CD20
SSC-A
PCDHγ
B
Gated on live CD3+
Donor A surface only
Donor A surface+secondary
Donor B surface only
Donor B surface+secondary
PCDHγ+
0
0
0
0.011
SSC-A
PCDHγ
C
Cohesin Chromatin Regulation Pathway
cAMP−mediated signaling
G−Protein Coupled Receptor Signaling
NF1 RAS Signaling Pathway
Degradation of the extracellular matrix
−log(p−value)
D
APH1A
APH1B
NCSTN
PSEN1
PSEN2
PSENEN
log2(TPM+1)
Tconv_naive
Tscm
Th2
Th1
Th17
Th1_17
cTfh
Treg_naive
Treg_mem
Temra

**Sup. Figure 9**

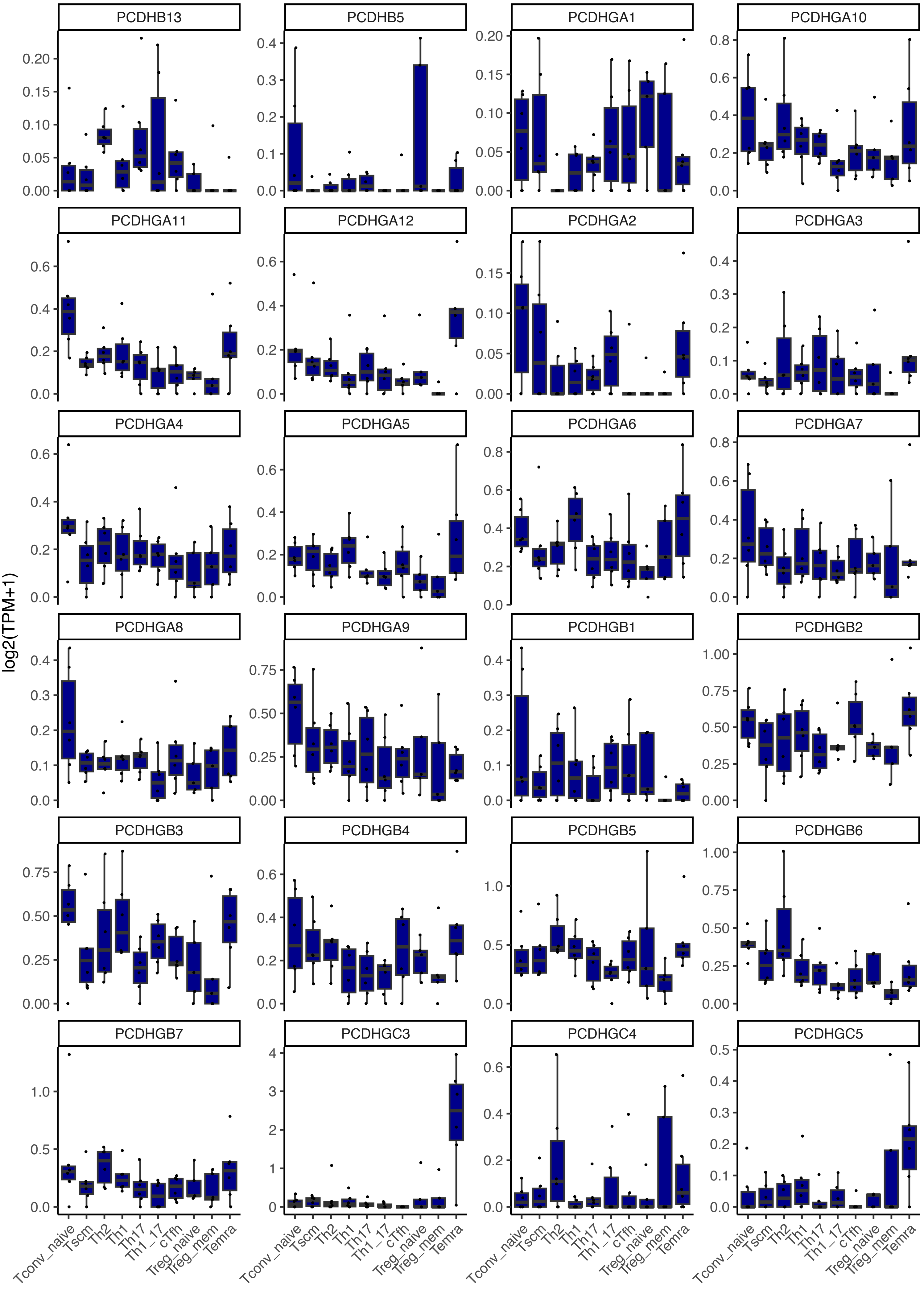

PCDHB13
PCDHB5
PCDHGA1
PCDHGA10
PCDHGA11
PCDHGA12
PCDHGA2
PCDHGA3
PCDHGA4
PCDHGA5
PCDHGA6
PCDHGA7
PCDHGA8
PCDHGA9
PCDHGB1
PCDHGB2
PCDHGB3
PCDHGB4
PCDHGB5
PCDHGB6
PCDHGB7
PCDHGC3
PCDHGC4
PCDHGC5
log2(TPM+1)
Tconv_naive
Tscm
Th2
Th1
Th17
Th1_17
cTfh
Treg_naive
Treg_mem
Temra
