## Supplementary figures and tables for "Epigenomic profiling of cerebrospinal fluid cells identifies immune regulatory alterations and implicates protocadherins in multiple sclerosis": Sup.Table1-3.docx

Table 1. Donor characteristics for the deep PBAT cohort

| Sample ID | Sex | Age | Diagnosis |
| --- | --- | --- | --- |
| 13-081 | M | 28 | NINDC |
| 13-100 | F | 37 | MS-rem |
| 13-101 | F | 19 | MS-rel |
| 13-138 | M | 33 | NINDC |
| 13-143 | F | 37 | NINDC |
| 14-013 | F | 21 | NINDC |
| 14-016 | M | 50 | MS-rel |
| 14-019 | F | 48 | MS-rel |
| 14-020 | F | 34 | MS-rem |
| 14-089 | F | 51 | NINDC |

Table 2. Independent donor characteristics for the shallow PBAT cohort

| Sample ID | Sex | Age | Diagnosis |
| --- | --- | --- | --- |
| 13-094 | F | 19 | MS-rel |
| 13-116 | M | 29 | MS-rel |
| 13-125 | F | 30 | MS-rel |
| 13-151 | F | 48 | MS-rel |
| 13-177 | F | 46 | MS-rel |
| 13-183 | F | 40 | MS-rel |
| 14-037 | F | 48 | MS-rel |
| 14-059 | F | 50 | MS-rel |
| 14-074 | F | 32 | MS-rel |
| 13-085 | F | 45 | MS-rem |
| 13-113 | F | 33 | MS-rem |
| 13-120 | F | 18 | MS-rem |
| 13-155 | F | 34 | MS-rem |
| 13-164 | F | 41 | MS-rem |
| 13-175 | F | 42 | MS-rem |
| 14-034 | F | 38 | MS-rem |
| 13-082 | F | 27 | MS-rem |
| 13-098 | F | 41 | MS-rem |
| 14-083 | F | 36 | NINDC |
| 13-091 | F | 21 | NINDC |
| 13-166 | M | 41 | NINDC |
| 14-007 | M | 26 | NINDC |
| 14-010 | M | 53 | NINDC |
| 14-077 | F | 61 | NINDC |
| 13-118 | M | 49 | NINDC |
| 13-123 | F | 21 | INDC |
| 13-192 | F | 40 | INDC |
| 14-018 | F | 50 | INDC |
| 14-023 | F | 28 | INDC |
| 14-070 | F | 37 | INDC |

Table 3. Overview of RNA-seq samples

| Sample ID | Group | Sequence |
| --- | --- | --- |
| S101 | NINDC | deep |
| S102 | RRMS | shallow |
| S103 | RRMS | shallow |
| S104 | NINDC | shallow |
| S105 | RRMS | shallow |
| S106 | RRMS | deep |
| S107 | NINDC | shallow |
| S108 | RRMS | shallow |
| S110 | NINDC | shallow |
| S111 | RRMS | shallow |
| S112 | NINDC | shallow |
| S113 | RRMS | deep |
| S114 | RRMS | deep |
| S115 | RRMS | deep |
| S116 | RRMS | shallow |
| S117 | NINDC | shallow |
| S13-183 | RRMS | shallow |
